# A TIR-1/SARM1 phase transition underlies p38 immune pathway activation in the *C. elegans* intestine

**DOI:** 10.1101/2021.08.05.455128

**Authors:** Nicholas D. Peterson, Janneke D. Icso, J. Elizabeth Salisbury, Tomás Rodríguez, Paul R. Thompson, Read Pukkila-Worley

## Abstract

Intracellular signaling regulators can be concentrated into membrane-free, higher-ordered protein assemblies to initiate protective responses during stress — a process known as phase transition. Here, we show that a phase transition of the *Caenorhabditis elegans* Toll/interleukin-1 receptor domain protein (TIR-1), an NAD^+^ glycohydrolase homologous to mammalian sterile alpha and TIR motif-containing 1 (SARM1), underlies p38 PMK-1 immune pathway activation in *C. elegans* intestinal epithelial cells. Through visualization of fluorescently labeled TIR-1/SARM1 protein, we demonstrate for the first time that physiologic stresses, both pathogen and non-pathogen, induce multimerization of TIR-1/SARM1 into visible puncta within intestinal epithelial cells. *In vitro* enzyme kinetic analyses revealed that, like mammalian SARM1, the NAD^+^ glycohydrolase activity of *C. elegans* TIR-1 is dramatically potentiated by protein oligomerization and a phase transition. Accordingly, *C. elegans* with genetic mutations that specifically block either multimerization or the NAD^+^ glycohydrolase activity of TIR-1/SARM1 fail to induce p38 PMK phosphorylation, are unable to increase immune effector expression, and are dramatically susceptible to bacterial infection. Finally, we demonstrate that the TIR-1/SARM1 phase transition is modified by dietary cholesterol, revealing a new adaptive response that allows a metazoan host to anticipate pathogen threats during micronutrient deprivation, a time of relative susceptibility to infection. When cholesterol is limited, TIR-1/SARM1 oligomerizes into puncta in intestinal epithelial cells and engages its NAD^+^ glycohydrolase activity, which increases p38 PMK-1 phosphorylation, and primes immune effector induction in a manner that promotes pathogen clearance from the intestine during a subsequent infection. Thus, a phase transition of TIR-1/SARM1 as a prerequisite for its NAD^+^ glycohydrolase activity is strongly conserved across millions of years of evolution and is essential for diverse physiological processes in multiple cell types.

## INTRODUCTION

The p38 mitogen-activated protein kinase (MAPK) pathway is a key regulator of stress responses and innate immune defenses in metazoans. The *C. elegans* p38 homolog PMK-1 is part of a classic MAPK signaling cascade that is activated by the MAPKKK NSY-1 and MAPKK SEK-1, which are the nematode homologs of mammalian ASK and MKK3/6, respectively [1]. *C. elegans* Toll/interleukin-1 receptor domain protein (TIR-1), an NAD^+^ glycohydrolase homologous to mammalian sterile alpha and TIR motif-containing 1 (SARM1), acts upstream of NSY-1 to control p38 PMK-1 activation [2, 3]. As in mammals, the *C. elegans* p38 PMK-1 pathway regulates the expression of secreted innate immune effectors and is required for survival during pathogen infection [1, 4–6]. However, the mechanisms that activate the NSY-1/SEK-1/p38 PMK-1 signaling cassette in *C. elegans* intestinal epithelial cells are poorly defined.

Intracellular signaling regulators can be compartmentalized in membrane-less, higher-ordered protein assemblies with liquid droplet-like properties [7–13]. Cytoplasmic de-mixing or phase transition of proteins in this manner concentrates signaling regulators to facilitate rapid and specific activation of protective defenses during stress [7, 8, 14]. Here, we show that a phase transition and NAD^+^ glycohydrolase activity of TIR-1/SARM1 is required for p38 PMK-1 immune pathway activation in *C. elegans* intestinal epithelial cells. By promoting aggregation of TIR-1/SARM1 with macromolecular crowding agents *in vitro*, we demonstrate that a phase transition is required for the catalytic activity of TIR-1/SARM1. Accordingly, TIR-1/SARM1 containing mutations that either specifically prevent the phase transition or impair NAD^+^ hydrolysis show decreased enzymatic activity *in vitro*. *C. elegans* carrying these same mutations, edited into the genome using CRISPR/Cas9, fail to induce p38 PMK phosphorylation, are unable to upregulate immune effector expression, and have enhanced susceptibility to bacterial infection. Importantly, we labeled the TIR-1/SARM1 protein with a fluorescent tag at its genomic locus and demonstrated for the first time that TIR-1/SARM1 oligomerizes into visible puncta within the intestine in response to physiologic stimuli, including both a pathogen and non-pathogen stress.

In a contemporaneous study, Loring et al. demonstrated that human SARM1 aggregates and undergoes a phase transition to potentiate its intrinsic NAD^+^ glycohydrolase activity [15]. These authors studied SARM1 in the context of neuronal degeneration as it was previously demonstrated that loss of mammalian SARM1 protects against axonal degeneration following neuronal injury [16–18]. Treatment with non-physiologic concentrations of citrate, a molecule that induces protein aggregation, led to oligomerization of *C. elegans* TIR-1 in neurons, which was correlated with enhanced axonal degeneration [15]. We demonstrate here that *C. elegans* TIR-1/SARM1 oligomerizes to form puncta in response to physiological stresses, providing an important characterization of the mechanism inferred by Loring, et al., and establish that the TIR-1/SARM1 phase transition occurs in non-neuronal tissues in the regulation of intestinal immunity. Thus, a phase transition of TIR-1/SARM1 as a prerequisite for its NAD^+^ glycohydrolase activity is strongly conserved, occurs in multiple cell types, and is essential for diverse physiological processes, including intestinal immune regulation and axonal degeneration.

Finally, we report that a non-pathogen stress can induce TIR-1/SARM1 oligomerization and p38 PMK-1 pathway activation as part of a mechanism to preempt pathogen attack during a time of relative vulnerability to infection. *C. elegans* lack the ability to synthesize cholesterol *de novo* and must acquire dietary sterols from its environment to support multiple aspects of cellular physiology, including development, fecundity, lifespan, and resistance against pathogen infection [19–25]. Some, but not all, *C. elegans* larvae that encounter sterol-scarce environments enter an alternative developmental program, called dauer diapause, to promote animal survival [26–30]. Here, we show that *C. elegans* that do not enter dauer diapause in an environment devoid of dietary sterols adapt by promoting oligomerization of TIR-1/SARM1 to activate the p38 PMK-1 innate immune pathway. Otarigho et al. previously demonstrated that dietary cholesterol is required for *C. elegans* to survive infection with the bacterial pathogen *Pseudomonas aeruginosa* and argued that cholesterol is necessary for the induction of some innate immune response pathways in nematodes [24]. Surprisingly, we found in multiple genome-wide transcriptome profiling experiments that *C. elegans* in a low cholesterol environment robustly activate, rather than repress, the transcription of a suite of genes, the majority of which are innate immune effectors that are downstream of the p38 PMK-1 pathway. We confirmed in genetic epistasis and western blot experiments that low cholesterol activates the TIR-1/NSY-1/SEK-1/p38 PMK-1 pathway in the absence of infection. Importantly, we show that priming p38 pathway activation in this manner augments immune effector expression during a subsequent bacterial infection and promotes pathogen clearance from the intestine. Thus, activation of the p38 PMK-1 pathway during conditions of low cholesterol availability is an adaptive response that allows nematodes to anticipate pathogen threats under conditions of essential metabolite scarcity.

## RESULTS

### Cholesterol scarcity activates intestinal innate immune defenses

*C. elegans* are sterol auxotrophs and require dietary sterols for development, lifespan, fecundity, and resistance to pathogen infection [19–25]. As such, 5 μg/mL of cholesterol is a standard additive in *C. elegans* laboratory growth medium [31]. We found that *C. elegans* grown in the absence of cholesterol supplementation activated GFP-based transcriptional reporters for two putative immune effector genes, T24B8.5p::*gfp* and *irg-5*p::*gfp* (Figs. 1A and B). T24B8.5 and *irg-5* are expressed in the intestine, induced during infection with multiple pathogens, including *P. aeruginosa,* and controlled by the p38 PMK-1 innate immune pathway [4, 32, 33]. qRT-PCR studies confirmed that *C. elegans* in a low cholesterol environment upregulate T24B8.5 and *irg-5,* as well as other innate immune effector genes (*irg-4* and K08D8.4) (Fig. 1C). These data suggest that host defense pathways are activated in the absence of pathogen infection when environmental sterols are scarce.

**Figure 1.**
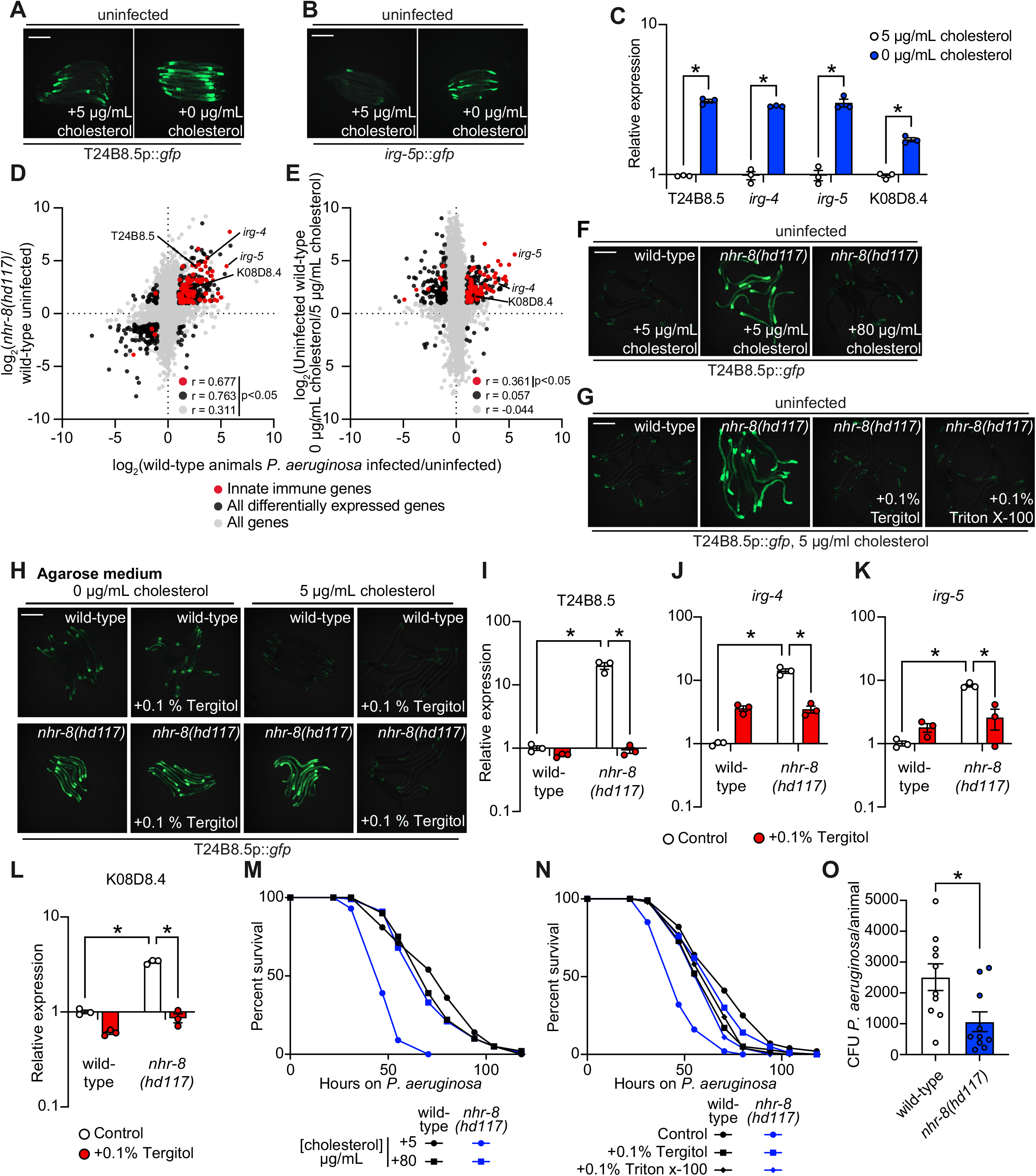
Cholesterol scarcity activates intestinal innate immune defenses. Images of T24B8.5p::*gfp* **(A)** and *irg-5*p::*GFP* **(B)** transcriptional immune reporters in wild-type animals growing on standard nematode growth media (+5 μg/mL cholesterol) and in the absence of supplemented cholesterol (+0 μg/mL cholesterol). **(C)** qRT-PCR data of the indicated innate immune effector genes in wild-type *C. elegans* growing in the presence (+5 μg/mL) and absence (+0 μg/mL) of supplemented cholesterol. *equals p<0.05 (unpaired t-test). **(D and E)** Data from mRNA-seq experiments comparing genes differentially regulated in uninfected *nhr-8(hd117)* mutants versus wild-type animals **(D)** or uninfected wild-type animals grown in the absence (0 μg/mL) versus presence (5 μg/mL) of supplemental cholesterol **(E)** (y-axis) are compared with genes differentially expressed in wild-type animals during *P. aeruginosa* infection (x-axis). All genes are shown in gray. Genes that are differentially expressed in both datasets are shown in black (Fold change >2, q<0.01). Genes that are annotated as innate immune genes are shown in red. The location of the representative genes T24B8.5, *irg-*5, irg*-4*, and K08D8.4, whose expression is examined throughout this manuscript are shown. See also Tables S1A-C. **(F, G, H)** Images of T24B8.5p::*gfp* animals of the indicted genotypes, grown under the indicated conditions are shown. **(H)** *C. elegans* were grown on media solidified with agarose, rather than agar. **(I, J, K, L)** qRT-PCR data of the indicated genes in wild-type and *nhr-8(hd117)* mutant animals grown on standard nematode growth media (+5 μg/mL cholesterol) in the presence or absence of 0.1% Tergitol, as indicated. For the qRT-PCR studies in Figs. 1C, 1I, 1J, 1K, and 1L, data are the average of three independent biological replicates, each normalized to a control gene with error bars representing SEM and are presented as the value relative to the average expression from all replicates of the indicated gene in wild-type animals on standard nematode growth media (+5 μg/mL cholesterol). *equals p<0.05 (two-way ANOVA with Tukey multiple comparison testing). **(M, N)** *C. elegans* pathogenesis assay with *P. aeruginosa* and *C. elegans* of indicated genotypes at the L4 larval stage is shown and exposed to the indicated conditions. Data are representative of three trials. The Kaplan-Meier method was used to estimate the survival curves for each group, and the log rank test was used for all statistical comparisons. Sample sizes, mean lifespan and p-values for all trials are shown in Table S2. **(O)** *P. aeruginosa,* isolated from the intestines of animals with the indicated genotypes, were quantified after 24 hours of bacterial infection. Data are colony forming units (CFU) of *P. aeruginosa* and are presented as the average of 10 separate biological replicates with each replicate containing 10-11 animals. *equals p<0.05 (unpaired t-test). Scale bars in all images equal 200 μm. See also Fig. S1.

The nuclear hormone receptor, NHR-8, a homolog of mammalian liver X receptor (LXR) and pregnane X receptor (PXR), is required for the transport, distribution, and metabolism of cholesterol in *C. elegans* [34, 35]. Thus, *nhr-8* loss-of-function mutant strains can be used as genetic tools to study conditions of low sterol content. Two previously-characterized *nhr-8* null alleles are *nhr-8(hd117)*, which lacks the first exon [34] and *nhr-8(ok186)*, which is missing most of the ligand-binding domain [35]. Notably, the transcription profile of *nhr-8(hd117)* and *nhr-8(ok186)* animals mimics that of wild-type *C. elegans* infected with the bacterial pathogen *P. aeruginosa* (Figs. 1D, S1A, Table S1A, and Table S1B). The correlation between the transcriptional signatures of either the *nhr-8(hd117)* or the *nhr-8(ok186)* mutant with the genes that are changed in wild-type animals during pathogen infection was significant across all genes (r = 0.311 and r = 0.370, respectively). Of note, the correlation between these datasets is tighter when comparing only the differentially expressed genes (r = 0.763 and r = 0.849, respectively) and only genes that are also involved in innate immunity (r = 0.677 and r = 0.703, respectively) (Figs. 1D and S1A). Among the immune effectors that are upregulated in both the *nhr-8(hd117)* and *nhr-8(ok186)* mutants, and in wild type animals infected with *P. aeruginosa,* are T24B8.5, *irg-4*, *irg-5*, and K08D8.4; the same genes whose transcription are also induced by cholesterol deprivation (Figs. 1C, 1D and S1A).

To validate these findings, we analyzed publicly available RNA-seq data that compared gene expression changes in wild-type *C. elegans* grown in media lacking supplemented cholesterol versus animals grown under standard culture conditions with 5 μg/mL of cholesterol [24]. Consistent with our transcriptome profiling experiments of *nhr-8* loss-of-function mutants, we found that innate immune effectors were strongly enriched among genes that were transcriptionally activated during cholesterol deprivation (Fig. 1E). Specifically, the expression of innate immune effectors was significantly correlated in wild-type animals infected with *P. aeruginosa* and in wild-type nematodes starved for cholesterol (r = 0.361) (Fig. 1E). Immune effectors that were induced during cholesterol starvation in qRT-PCR studies (*irg-4*, *irg-5* and K08D8.4, but not T24B8.5) were again found among these differentially regulated genes, confirming the integrity of our RNA-seq analysis (Fig. 1C). In summary, cholesterol-starved, wild-type *C. elegans* and two different *nhr-8* loss-of-function mutants induce the transcription of innate immune defenses.

Two different supplementation experiments using the T24B8.5p::*gfp* immune reporter demonstrated that immune effector activation in *nhr-8(hd117)* mutants is due to sterol deficiency in these animals. First, supplementation of exogenous cholesterol at an increased concentration (80 μg/mL) fully suppressed T24B8.5p::*gfp* activation in the *nhr-8(hd117)* mutant (Fig. 1F). Second, supplementation with the non-ionic detergents Tergitol or Triton X-100, which solubilize hydrophobic, amphipathic compounds, including sterols, suppressed T24B8.5p::*gfp* activation in *nhr-8(hd117)* animals in a manner that was dependent on the presence of added cholesterol in the growth media (Figs. 1G, 1H and S1B). Nematode growth media solidified with agarose, rather than agar, contains markedly fewer contaminating sterols [20, 28, 34]. Importantly, on agarose, the addition of Tergitol to the growth media also suppressed T24B8.5p::*gfp* activation in the *nhr-8(hd117)* mutant, but only in the presence of 5 μg/mL of cholesterol (Fig. 1H). Notably, this effect was dose-dependent (Fig. S1B). These data establish that solubilization of cholesterol by Tergitol in standard nematode growth media suppresses immune activation in the *nhr-8(hd117)* mutant background. We used qRT-PCR to confirm these findings – Tergitol suppressed activation of T24B8.5, as well as the immune effectors *irg-4*, *irg-5* and K08D8.4 in the *nhr-8(hd117)* mutant (Figs. 1I-L). Of note, activation of immune defenses in *nhr-8(hd117)* animals is specific to cholesterol deprivation in this genetic background, as supplementation with individual unsaturated, mono- and polyunsaturated fatty acids, which are also mis-regulated in the *nhr-8(hd117)* mutant background [34], failed to suppress activation of T24B8.5p::*gfp* (Fig. S1C). Together, these data confirm that cholesterol deprivation activates immune effector transcription independent of bacterial infection.

It has been previously shown that *C. elegans* raised on media without supplemented cholesterol are hypersusceptible to killing by *P. aeruginosa*, as are *nhr-8(hd117)* mutants on media with standard cholesterol supplementation (5 μg/mL) [24]. Importantly, we found that high doses of cholesterol (80 μg/mL) rescued the enhanced susceptibility of *nhr-8(hd117)* mutants to *P. aeruginosa* infection (Fig. 1M). We confirmed this finding in an orthologous approach. Two detergents (Tergitol or Triton X-100), which solubilize cholesterol in nematode growth media, also restored wild-type pathogen resistance to the *nhr-8(hd117)* mutant (Fig. 1N), consistent with their ability to modulate the expression of innate immune effectors (Fig. 1G-L). Thus, cholesterol is required for pathogen resistance in *C. elegans*.

Importantly, we found that the robust transcriptional induction of immune effectors observed in the *nhr-8(hd117)* mutant promotes clearance of *P. aeruginosa* from the intestine (Fig. 1O). Thus, even though *nhr-8(hd117)* mutant animals are more susceptible to pathogen killing, they are mounting effective anti-pathogen defenses toward an ingested pathogen (Fig. 1O). These data suggest that *C. elegans* activates host immune defenses when environmental sterols are limited to preempt pathogen attack during a period of relative susceptibility to bacterial infection.

### Cholesterol scarcity activates the p38 PMK-1 innate immune pathway

Additional analyses of the transcriptome profiling data revealed that targets of the p38 PMK-1 innate immune pathway were strongly enriched among the genes induced in wild-type animals during cholesterol deprivation (Fig. 2A). Consistent with this observation, the levels of active, phosphorylated PMK-1 were higher in wild-type animals grown on media without supplemented cholesterol than in animals grown under standard culture conditions (Figs. 2B and 2C). These data demonstrate that cholesterol deprivation activates p38 PMK-1.

**Figure 2.**
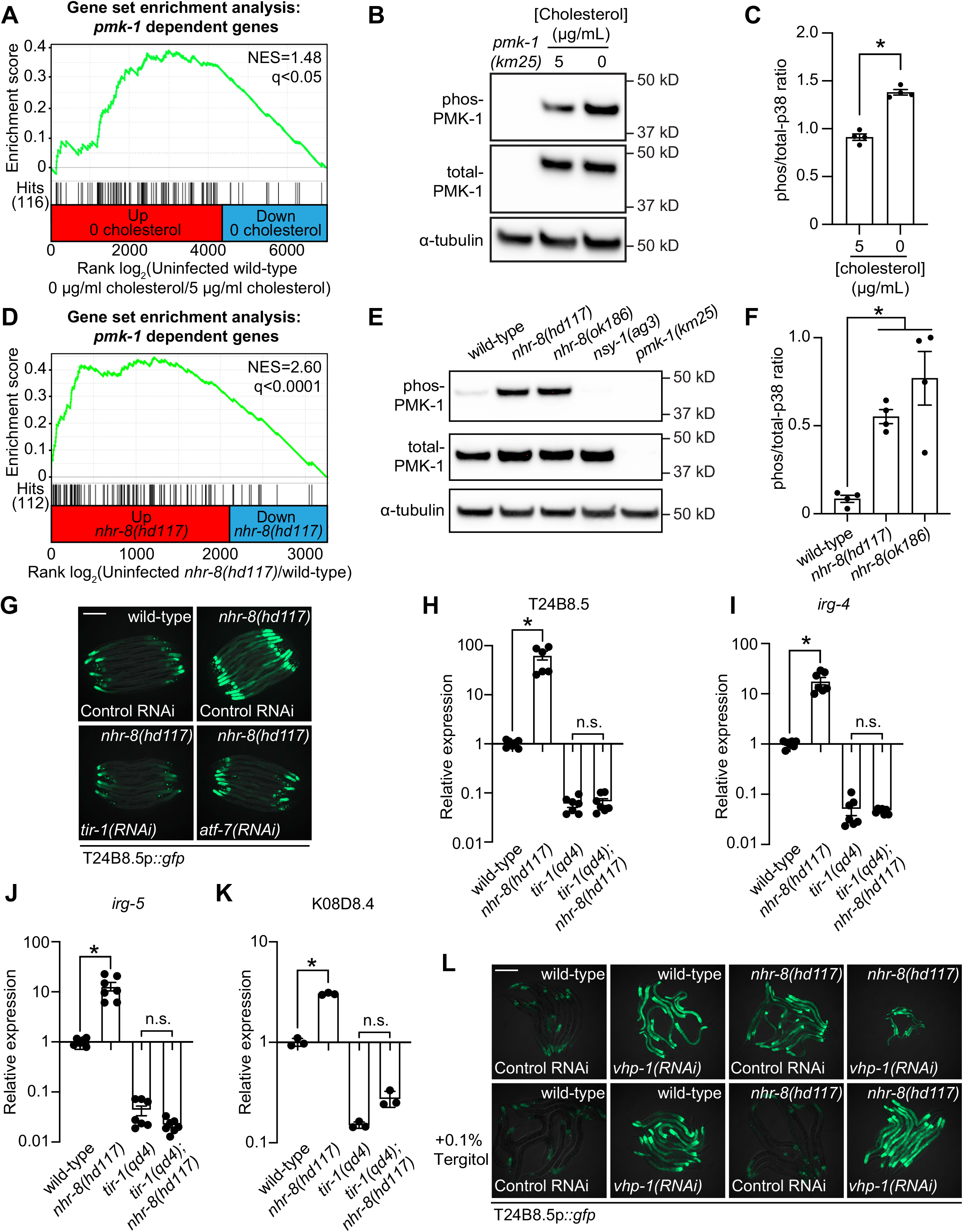
Cholesterol scarcity activates the p38 PMK-1 innate immune pathway. **(A)** Gene set enrichment analysis (GSEA) of p38 PMK-1 targets in the 0 μg/mL cholesterol mRNA-seq experiment. Fold change in expression of significantly differentially expressed genes (fold-change >2 and q<0.01) in uninfected animals grown in the absence (0 μg/mL) versus presence (5 μg/mL) of supplemental cholesterol are ranked from higher expression (red) to lower expression (blue). Normalized enrichment score (NES) and q-value are indicated. p38 PMK-1 targets found in the transcriptional profile are indicated by hit number (116) in the left margin and black lines. **(B)** An immunoblot analysis of lysates from wild-type *C. elegans* grown on standard nematode growth media in the presence (+5 μg/mL cholesterol) and in the absence (+0 μg/mL cholesterol) of supplemented cholesterol using antibodies that recognize the doubly phosphorylated TGY motif of PMK-1 (phos-PMK-1), total PMK-1 protein (total PMK-1) and tubulin (α-tubulin) is shown. PMK-1 is a 43.9 kDa protein and tubulin is a 50 kDa protein. **(C)** The band intensities of four biological replicates of the Western blot shown in Fig. 2A were quantified. *equals p<0.05 (unpaired t-test) **(D)** GSEA of p38 PMK-1 targets in the *nhr-8* mRNA-seq experiment as described in Fig. 2A. **(E, F)** Western blot experiment and quantification of four biological replicate experiments as described in Fig. 2B and Fig. 2C with the strains of the indicated genotypes. In Fig. 2B and Fig. 2E, *pmk-1(km25)* and *nsy-1(ag3)* and loss-of-function mutants are the controls, which confirm specificity of phos-PMK-1 and total PMK-1 probing. *equals p<0.05 (one-way ANOVA with Dunnett multiple comparison testing). **(G)** Images of T24B8.5p::*gfp* transcriptional immune reporter animals in wild-type animals and *nhr-8(hd117)* mutants growing on control RNAi, *tir-1(RNAi)* or *atf-7(RNAi)* bacteria, as indicated. **(H, I, J, K)** qRT-PCR data of the indicated genes in the indicated mutant animals grown on standard nematode growth media (+5 μg/mL cholesterol). Data are the average of three to seven independent replicates, each normalized to a control gene with error bars representing SEM and are presented as the value relative to the average expression from all replicates of the indicated gene in wild-type animals. *equals p<0.05 (one-way ANOVA with Dunnett multiple comparison testing) **(L)** Images of T24B8.5p::*gfp* transcriptional immune reporter animals of the indicated genotypes growing in the presence or absence of 0.1% Tergitol. Scale bars in all images equal 200 μm. See also Fig. S2.

The p38 PMK-1 pathway is also activated in *nhr-8* loss-of-function mutants. A gene set enrichment analysis of the *nhr-8(hd117)* and *nhr-8(ok186)* transcriptome profiling experiments revealed strong enrichment of p38 PMK-1 pathway targets among the genes upregulated in each *nhr-8* mutant (Figs. 2D and S2A). Consistent with these data, the *nhr-8(hd117)* and *nhr-8(ok186)* mutants had an increased ratio of phosphorylated PMK-1 relative to total PMK-1 compared to wild-type controls (Figs. 2E and 2F). In addition, RNAi-mediated knockdown of *tir-1*, the most upstream component of the p38 signaling cassette [2, 3], fully suppressed hyperactivation of T24B8.5p::*gfp* in *nhr-8(hd117)* animals (Fig. 2G). In addition, the *tir-1(qd4)* loss-of-function mutation completely suppressed the induction of T24B8.5, *irg-4*, *irg-5*, and K08D8.4 in the *nhr-8(hd117)* background (Figs. 2H-K). The transcription factors ATF-7 and SKN-1 link PMK-1 activity to its transcriptional outputs [36, 37]. In *nhr-8(hd117)* animals, knockdown of *atf-7*, but not *skn-1,* abrogated T24B8.5p::*gfp* activation (Figs. 2G and S2B). Finally, to further support our observation that cholesterol scarcity induces immune defenses upstream of p38 PMK-1, we used the MAPK phosphatase *vhp-1*, a negative regulator of PMK-1 [38]. Solubilization of cholesterol with Tergitol was unable to suppress activation of T24B8.5p::*gfp* induced by knockdown of *vhp-1* (Fig. 2L). Thus, the TIR-1-p38 PMK-1-ATF-7 signaling axis is activated in the *nhr-8* mutant background.

The c-JUN N-terminal kinase MAPK homolog *kgb-1*, the insulin signaling pathway forkhead box O family (FOXO) transcription factor, *daf-16*, and the G protein-coupled receptor (GPCR), *fshr-1*, each function in parallel to the p38 PMK-1 pathway to regulate immune and stress responses in *C. elegans* [4, 38–40]. However, knockdown of each of these genes in *nhr-8(hd117)* animals failed to suppress T24B8.5p::*gfp* activation (Fig. S2B) and RNAi-mediated knockdown of *nhr-8* did not induce nuclear localization of DAF-16::GFP (Fig. S2C). Thus, cholesterol scarcity induces *C. elegans* innate immune responses through specific activation of p38 PMK-1 immune pathway signaling.

### Activation of the p38 PMK-1 pathway by cholesterol deprivation requires TIR-1 oligomerization and NAD^+^ glycohydrolase activity

*C. elegans* TIR-1, which is homologous to mammalian SARM1, functions upstream of the MAPKKK NSY-1; MAPKK SEK-1; MAPK p38 PMK-1 signaling cassette in the intestine to control innate immune defenses [2, 41]. Accordingly, *tir-1* loss-of-function mutants are more susceptible to killing by *P. aeruginosa* than wild-type animals [2]. The TIR-1 protein has three characterized domains: a Heat/Armadillo repeat domain, a sterile alpha motif (SAM) domain, and a Toll-interleukin receptor (TIR) domain [41] (Fig. 3A). *C. elegans* TIR-1 oligomerizes *in vitro* through interactions of its SAM domains [42]. In addition, oligomerization of SARM1 is essential for its function in plants and mice [42, 43]. As such, we evaluated whether the oligomerization of *C. elegans* TIR-1 is required to activate the p38 PMK-1 immune pathway in the intestine. We used CRISPR-Cas9 to delete both SAM domains in *tir-1* (*tir-1*^ΔSAM^) (Fig. 3A). We also generated point mutants in two residues within the *C. elegans* TIR domain that are important for the self-association and activity of mammalian SARM1 (*C. elegans tir-1*^G747P^ and *tir-1*^H833A^) [42] (Fig. 3A). The *C. elegans tir-1*^ΔSAM^*, tir-1*^G747P^ and *tir-1*^H833A^ mutants prevented activation of the p38 PMK-1-dependent immune reporter T24B8.5p::*gfp* in animals infected with *P. aeruginosa* (Fig. 3B), in *nhr-8(RNAi)* animals (Fig. 3C), and during cholesterol deprivation (Fig. 3D). Consistent with these data, the *tir-1*^ΔSAM^*, tir-1*^G747P^ and *tir-1*^H833A^ mutants have reduced levels of active, phosphorylated p38 PMK-1, equivalent to the *tir-1(qd4)* null allele [32] (Figs. 3E and 3F). Additionally, these mutants abrogate the induction of the T24B8.5 (Fig. 3G) and *irg-4* (Fig. S3A) immune effectors in *nhr-8(RNAi)* animals and are each markedly hypersusceptible to *P. aeruginosa* infection (Fig. 3H). The TIR domain of *C. elegans*, TIR-1, and its mammalian homolog, SARM1, possess intrinsic NADase activity [42, 44, 45]. Importantly, oligomerization of mammalian SARM1 and *C. elegans* TIR-1 is required for maximal NADase activity *in vitro* [42]. The NADase activity in the TIR domain of mammalian SARM1 requires a putative catalytic glutamate [45]. We used CRISPR-Cas9 to mutate the homologous glutamate in *C. elegans tir-1* (*tir-1*^E788A^) and found that it was required for the immunostimulatory activity of *tir-1 – tir-1*^E788A^ mutants do not induce T24B8.5p::*gfp* following *P. aeruginosa* infection (Fig. 3B), knockdown of *nhr-8* (Fig. 3C) or during cholesterol scarcity (Fig. 3D). Likewise, the *tir-1*^E788A^ mutant animals had less active, phosphorylated p38 PMK-1 (Figs. 3E and 3F), completely suppressed the induction of T24B8.5 (Fig. 3G) and *irg-4* (Fig. S3A) and were more susceptible to *P. aeruginosa* infection (Fig. 3H). We confirmed that *tir-1*^E788A^, *tir-1*^ΔSAM^*, tir-1*^G747P^ and *tir-1*^H833A^ mutants are translated and not degraded by introducing a 3xFLAG tag at the C-terminus of each mutant using CRISPR-Cas9 (Figs. S3B and S3C). A western blot with an anti-FLAG antibody yielded a band of the expected size for each *tir-1* mutant (Fig. S3B). In addition, the 3xFLAG-tagged wild-type TIR-1 expressed T24B8.5p::gfp normally, but the tagged mutant TIR-1 proteins did not (Fig. S3C). Collectively, these data provide the first genetic evidence that both the oligomerization of TIR-1 and its intrinsic NADase activity are required to activate the p38 PMK-1 innate immune pathway.

**Figure 3.**
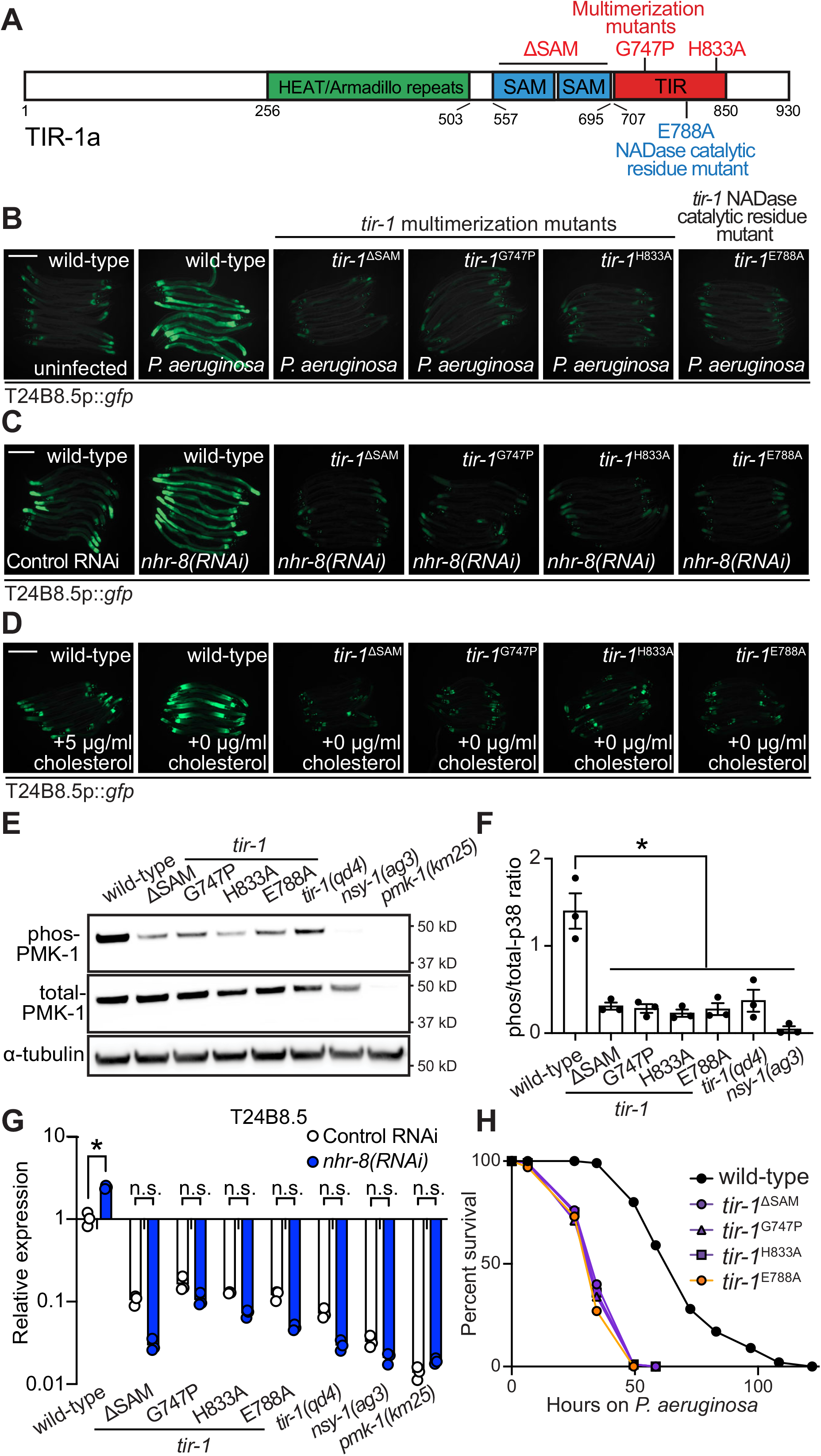
Activation of the p38 PMK-1 pathway by cholesterol deprivation requires TIR-1 oligomerization and NADase activity. **(A)** Model of *tir-1* showing the domains and the mutations that were introduced using CRISPR-Cas9. **(B, C, D)** Expression of the innate immune effector T24B8.5p::*gfp* in *tir-1* mutants with predicted defects in oligomerization (*tir-1*^ΔSAM^*, tir-1*^G747P^ and *tir-1*^H833A^) and NADase catalytic activity (*tir-1*^E788A^) during *P. aeruginosa* infection **(B)**, following *nhr-8(RNAi)*, and during cholesterol deprivation **(D)**. Scale bars in all images equal 200 μm. **(E)** Immunoblot analysis of lysates from the indicated genotypes probed with antibodies targeting the doubly phosphorylated TGY epitope in phosphorylated PMK-1 (phos-PMK-1), total PMK-1 protein (total PMK-1), and tubulin (α-tubulin). *nsy-1(ag3)* and *pmk-1(km25)* loss-of-function mutants are the controls, which confirm the specificity of the phospho-PMK-1 probing. **(F)** The band intensities of three biological replicates of the Western blot shown in **(E)** were quantified. *equals p<0.05 (one-way ANOVA with Dunnett multiple comparison testing). **(G)** qRT-PCR data of T24B8.5 in wild-type and mutant animals of the indicated genotypes grown on standard nematode growth media (+5 μg/mL cholesterol). Data are the average of three independent replicates, each normalized to a control gene with error bars representing SEM and are presented as the value relative to the average expression from all replicates in wild-type animals. *equals p<0.05 (two-way ANOVA with Tukey multiple comparison testing). (H) *C. elegans* pathogenesis assay with *P. aeruginosa* and *C. elegans* of indicated genotypes at the L4 larval stage are shown. Data are representative of three trials. The Kaplan-Meier method was used to estimate the survival curves for each group, and the log rank test was used for all statistical comparisons. Sample sizes, mean lifespan and p-values for all trials are shown in Table S2. See also Fig. S3.

### The NAD^+^ glycohydrolase activity of TIR-1/SARM1 is activated by a phase transition

To further characterize the mechanism of TIR-1/SARM1 activation, we recombinantly expressed and purified the TIR domain of the TIR-1 protein (called TIR) from *E. coli* and evaluated its NADase activity *in vitro* using an etheno-NAD^+^ (ε-NAD) activity assay, in which hydrolysis of the nicotinamide ring of ε-NAD leads to an increase in fluorescence. Interestingly, purified TIR only shows very modest NADase activity even at high protein concentrations (>15 μM) (Fig. 4A). Notably, the NADase activity of TIR increased parabolically with increasing TIR concentrations rather than linearly, suggesting that multimerization of TIR-1 potentiates its NADase activity (Fig. 4A).

**Figure 4.**
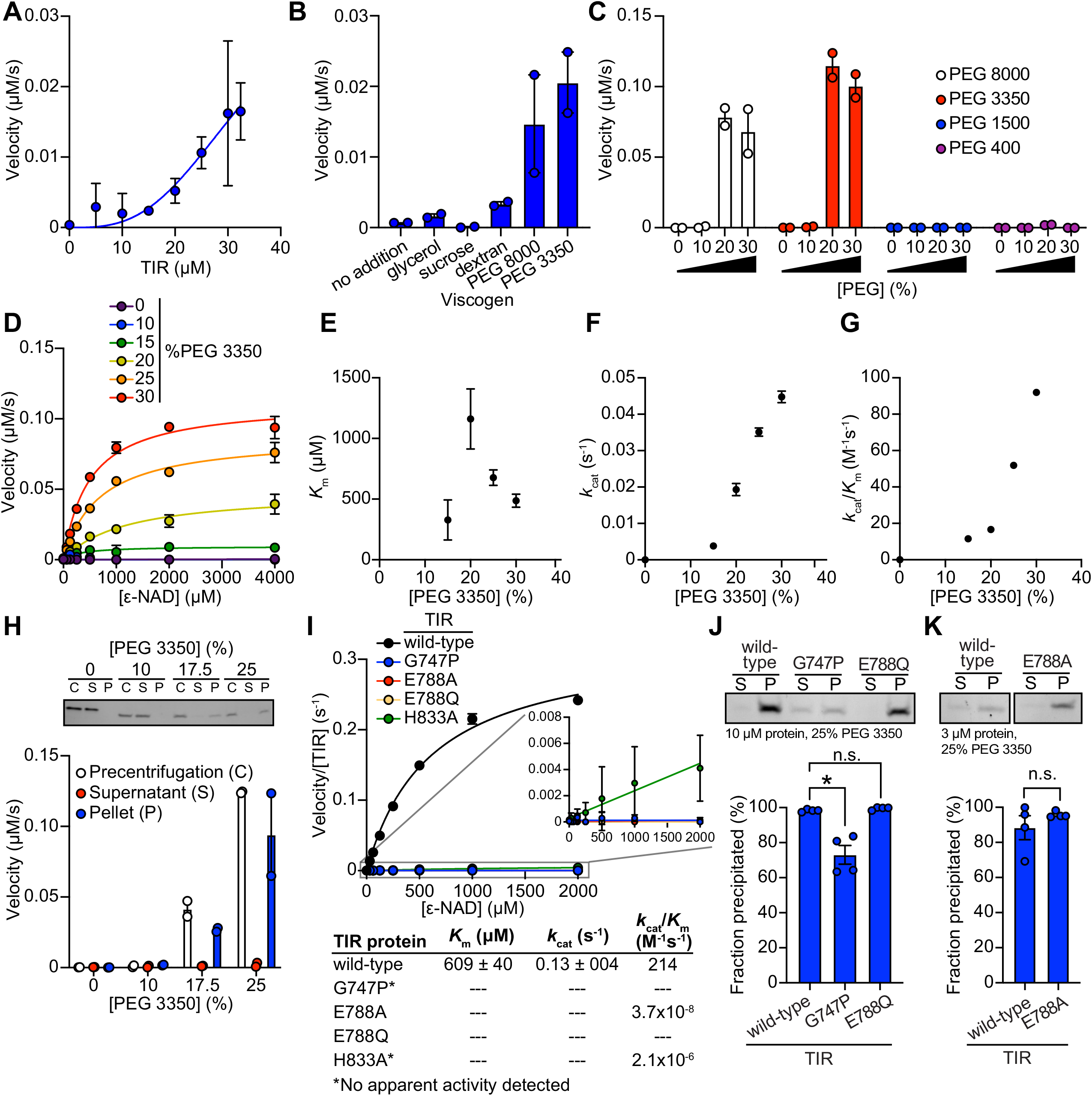
The NAD^+^ glycohydrolase activity of TIR-1/SARM1 is activated by a phase transition that requires TIR oligomerization. **(A)** NADase activity of purified TIR at increasing TIR protein concentrations is shown. Activity was assessed by incubating TIR protein with 1 mM ε-NAD and monitoring the rate at which the fluorescent product ε-ADPR was produced. Curve represents a nonlinear regression fit of the NADase activity data points (n=2). **(B)** NADase activity of 2.5 μM TIR incubated in the presence of 25% (w/v) of macro- (PEG 8000, PEG 3350 and dextran) and micro- (sucrose and glycerol) viscogens was assessed as described in Fig. 4A (n=2). **(C)** Dose dependency of macroviscogens on the NADase activity of TIR is shown. 2.5 μM TIR protein was incubated with the indicated PEG compound at concentrations from 0-30% (w/v). NADase activity was assessed as described in Fig. 4A (n=2). **(D)** Steady-state kinetic analysis of 2.5 µM TIR incubated in 0-30% (w/v) of PEG 3350 with the ε-NAD substrate provided at concentrations from 0-4000 μM was assessed as described in Fig. 4A. (n=2). From the steady-state kinetic analysis performed in Fig. 4D, *K*_m_ **(E)**, *k*_cat_ **(F)**, and *k*_cat_/*K*_m_ **(G)** were determined at each PEG 3350 concentration. **(H)** SDS-PAGE analysis of TIR protein fractions incubated with increasing concentrations of PEG 3350 before [precentrifugation (C)] and after centrifugation, the soluble (S) and pellet (P) protein fractions. NADase activity of TIR protein in each fraction and at each concentration of PEG 3350 was assessed, as described in Fig. 4A and is represented below the gel image (n=2, representative image shown). **(I)** Steady-state kinetic analysis of TIR wild-type, oligomerization mutant (TIR^G747P^, TIR^H833A^), and catalytic mutants (TIR^E788Q^ and TIR^E788A^) in 25% PEG 3350 assessed with 0-2000 μM ε-NAD was assessed as described in Fig. 4D. Kinetic parameters (*K*_m_, *k*_cat_ and *k*_cat_/*K*_m_) are shown in the table below the graph (n=3). **(J, K)** SDS-PAGE analysis of TIR: wild-type, oligomerization mutant (TIR^G747P^) and catalytic mutants (TIR^E788Q^ and TIR^E788A^), precipitation in the presence of 25% PEG 3350. Gel represents the soluble (S) and pellet (P) protein fractions of wild-type and mutant TIR following incubation with PEG 3350 and centrifugation. TIR^G747P^ and TIR^E788Q^ were assessed with 10 μM protein in Fig. 4J and TIR^E788A^ was assessed with 3 μM protein in Fig. 4K. Quantification of replicates represented below gel images (n=4, representative images shown). *equals p<0.05 by one-way ANOVA in Fig. 4J and unpaired t-test in Fig. 4K. See also Fig. S4.

Given that high concentrations of TIR are required to observe NADase activity, we hypothesized that molecular crowding might activate the enzyme and therefore assessed the effect of several macro- and microviscogens on TIR activity. Macroviscogens reduce the free volume available for protein movement and thus promote aggregation of protein complexes that are capable of self-association [46, 47]. Importantly, macroviscogens have minimal impact on the rate of diffusion of small molecules. By contrast, microviscogens, which are much smaller than most enzymes, affect the diffusion of substrates in solution and thus, the frequency at which enzymes encounter their substrate [47]. Interestingly, macroviscogens [polyethylene glycol (PEG) 3350 and PEG 8000], but not microviscogens (sucrose or glycerol), dramatically increased the NADase activity of TIR (Fig. 4B). PEGs 3350 and 8000 increase TIR activity in a concentration-dependent manner (Fig. 4C). These effects were most pronounced with higher molecular weight PEGs, as treatment with smaller molecular weight PEGs (e.g., PEG 1500 and PEG 400) or dextran did not increase the enzymatic activity of TIR (Fig. 4C). Crowding agents also increase the activity of the TIR domain of human SARM1, as well as plant TIR domains [15, 42], suggesting that the mechanism of TIR regulation is strongly conserved.

Treatment with macroviscogens enabled the characterization of the enzyme kinetics of TIR. Incubation with 25% PEG 3350 led to a roughly linear association between TIR concentration and NADase activity and markedly enhances TIR activity at each enzyme concentration (Fig. S4A). Consequently, lower amounts of enzyme (2.5 μM TIR) can be used to obtain robust kinetic data. Using these conditions, we determined the steady-state kinetic parameters (*k*_cat_, *K*_M_, and *k*_cat_/*K*_M_) with 2.5 μM TIR in the presence of increasing concentrations of PEG 3350 (Fig. 4D). The *K*_m_ of TIR increases and then decreases to level off at ∼500 μM of ε-NAD with increasing concentration of PEG 3350 (Fig. 4E). On the other hand, *k*_cat_ increases nearly linearly with increasing concentrations of PEG 3350 (Fig. 4F). In addition, the catalytic efficiency (*k*_cat_/*K*_m_) followed the same trend as *k*_cat_, indicating that increased TIR activity is due to increased substrate turnover by the enzyme and not tighter substrate binding (Fig. 4G).

We found that the NAD hydrolase activity of TIR is also activated by the precipitant sodium citrate, providing an orthologous method to characterize the enzymatic activity of the TIR domain (Figs. S4A and S4B). Notably, the response to increasing citrate concentration was switch-like, where a concentration of at least 250 mM sodium citrate was needed to observe enzyme activity (Figs. S4B and S4C). By contrast, activation with PEG 3350 was more dose-dependent (Figs. 4C and 4D). Nevertheless, the kinetic parameters obtained in the presence of citrate displayed similar trends to those found with PEG 3350: *K*_m_ decreased to level off at ∼400 µM of ε-NAD (Fig. S4D), *k*_cat_ increases with increasing sodium citrate concentration (Fig. S4E), and *k*_cat_/*K*_m_ follows the *k*_cat_ trends (Fig. S4F).

In the cytoplasm, high concentrations of macromolecules (proteins, nucleic acids, lipids, carbohydrates) cause molecular crowding and induce phase transitions of signaling proteins [48]. To determine if TIR undergoes a phase transition and whether its NADase activity correlates with the transition, we incubated purified TIR with different concentrations of PEG3350 or citrate, centrifuged the sample, and evaluated TIR NADase activity in the soluble and insoluble fractions (Figs. 4H and S4G). At low concentrations of both PEG and citrate, TIR protein was present in the supernatant. However, at high concentrations of both PEG and citrate, TIR was principally located in the pelleted (insoluble) fraction, as visualized by Coomassie staining on an SDS-PAGE gel (Figs. 4H and S4G). Importantly, robust NADase activity was observed in the pelleted fraction, and not in the supernatant of samples treated with high concentrations of PEG or citrate (Figs. 4H and S4G). Taken together, these data demonstrate that precipitation/aggregation of TIR is required to activate the intrinsic NADase activity of TIR-1/SARM1.

### TIR phase transition and NAD^+^ glycohydrolase activity requires TIR oligomerization

To determine whether TIR-1 oligomerization is also required for a phase transition and the NADase activity of TIR, we recombinantly expressed and purified the TIR domain of TIR-1, which contained mutations in residues required for oligomerization (TIR^G747P^ and TIR^H833A^) and NAD^+^ hydrolysis (TIR^E788A^ and TIR^E788Q^). Notably, TIR^G747P^ and TIR^E788Q^ mutants showed no apparent activity and TIR^E788A^ and TIR^H833A^ showed a >1×10^8^-fold decrease in catalytic efficiency (*k*_cat_/K_m_) compared to TIR^WT^ (Fig. 4I). Similarly, kinetic analysis of TIR oligomerization and catalytic mutants in 500 mM citrate revealed that all the mutants were catalytically dead, each showing no apparent activity (Fig. S4H).

To further characterize the TIR phase transition, we evaluated the precipitation capacity of TIR oligomerization and catalytic mutants in the presence of PEG 3350 and citrate. Consistent with our genetic data, TIR^G747P^, which contains a mutation that prevents oligomerization of TIR, was unable to precipitate as readily as TIR^WT^ (Fig. 4J). Compared to TIR^WT^, the TIR^G747P^ mutant in the presence of PEG 3550 demonstrated a 25% decrease in the amount of precipitated protein (Fig. 4J). We observed similar results in the presence of citrate, in which compared to TIR^WT^, the TIR^G747P^ mutant demonstrated a 22% decrease in precipitated protein compared to TIR^WT^ (Fig. S4I). Importantly, TIR proteins with two different mutations in the glutamate required for NAD^+^ catalysis, TIR^E788A^ and TIR^E788Q^, precipitated to a similar extent as TIR^WT^ (Figs. 4J, 4K, S4I, and S4J), but had minimal NADase activity (Figs. 4I and S4H). Collectively, these data demonstrate that a phase transition of TIR-1 is required for its NADase activity.

### A phase transition superactivates the intrinsic NAD^+^ glycohydrolase activity of TIR

During phase separations, macromolecules partition into distinct biochemical compartments characterized by a higher concentration dense phase and a lower concentration dilute phase [10, 11, 13]. The dense compartment can have either liquid-like properties in liquid-to-liquid phase separations or solid-like properties in liquid-to-solid phase transitions [11, 12, 14, 49]. To determine whether TIR undergoes a liquid-to-liquid or a liquid-to-solid phase transition, we assayed the enzyme activity of TIR in the presence of 1,6-hexanediol, an aliphatic alcohol that interferes with hydrophobic interactions prominent in liquid-to-liquid separations. Thus, 1,6-hexanediol disrupts liquid-like compartments, but not solid-like compartments [10, 13, 14]. In the presence of either PEG 3350 or citrate, we observed that 1,6-hexanediol decreases the NADase activity of TIR by <2-fold, yet the activity of 1,6-hexanediol-treated TIR remains two orders of magnitude higher than the activity of TIR without PEG 3550 or citrate addition (Fig. 5A). While significant, this modest effect suggests that the NADase activity of TIR is predominantly associated with a solid, rather than a liquid, state (Fig. 5A). Notably, in the absence of PEG 3350 or citrate, the low-level TIR NADase activity can be inhibited by 1,6-hexanediol (Fig. 5A), suggesting that this activity is driven by hydrophobic interactions. Importantly, 1,6-hexanediol does not interfere with TIR precipitation in the presence of PEG or citrate, regardless of whether it is added before or after precipitate formation (Figs. S5A and S5B).

**Figure 5.**
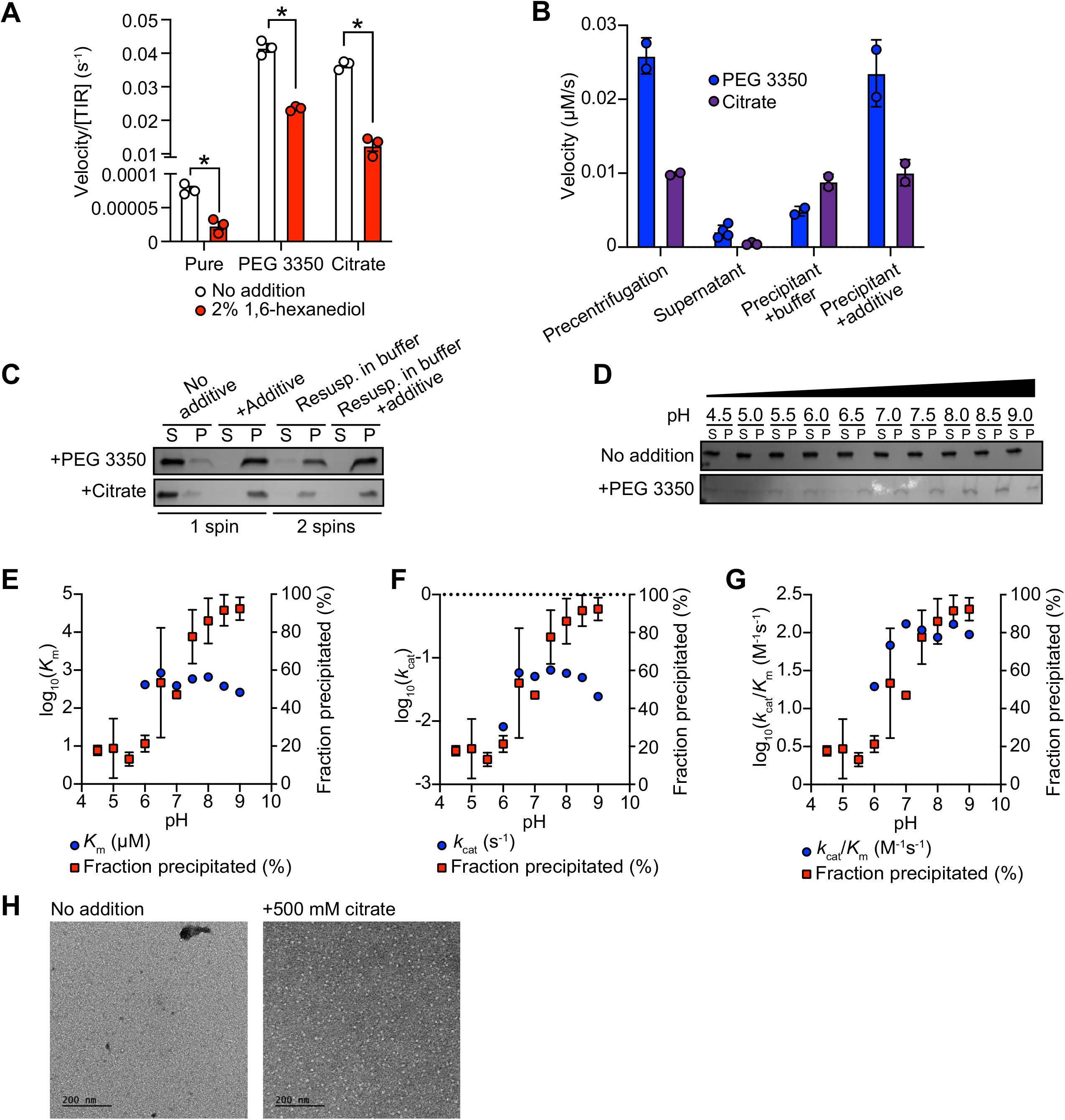
A phase transition superactivates the intrinsic NAD^+^ glycohydrolase of TIR. **(A)** Effect of 1,6-hexanediol on TIR NADase activity is shown. TIR protein was incubated in the presence or absence of either 25% PEG 3350 or 500 mM citrate and treated with either 0 or 2% 1,6-hexanediol. The NADase activity of TIR for each condition was assessed using the ε-NAD substrate assay (n=3). *equals *p*<0.05 (unpaired t-test). **(B)** The NADase activity of TIR protein incubated with either 25% PEG 3350 or 500 mM citrate before (precentrifugation, n=2) and after centrifugation, the supernatant (n=4) and precipitant (n=2) fractions. Precipitation fractions were resuspended in buffer alone or buffer containing 25% PEG 3350 or 500 mM citrate and NADase activity was assessed. **(C)** SDS-PAGE analysis of TIR protein in the soluble (S) and pellet (P) fractions following incubation with either PEG 3350 or citrate and centrifugation (1 spin). Pellet fractions were subsequently resuspended in only buffer or buffer containing 25% PEG 3350 or 500 mM citrate and centrifuged a second time to isolate the soluble (S) and pellet (P) fractions (n=2, representative image shown). **(D)** SDS-PAGE analysis of TIR in the soluble (S) and pellet (P) fractions incubated with or without 25% PEG 3350 at the indicated pH (n=2, representative image shown). TIR steady state kinetic parameters [*K*_m_ **(E)**, *k*_cat_ **(F)**, *k*_cat_/*K*_m_ **(G)**] is shown at the indicated pH (n=2). **(H)** Negative stain electron microscopy in either the absence or presence of 500 mM citrate (diameter of particles = 8.9 nm ± 1.2, n = 65). See also Fig. S5.

Next, we determined if this liquid-to-solid phase transition of TIR is reversible. The insoluble fraction following treatment of TIR with PEG 3350 or citrate was resuspended in either buffer alone or buffer plus the respective additive (PEG 3350 or citrate). If the phase transition is reversible, resuspension of precipitated TIR in buffer alone should disrupt enzymatic activity. However, if the phase transition is irreversible, activity should be detected when the pellet is resuspended in buffer alone. Notably, we observed some TIR enzymatic activity when the pellet was resuspended in buffer (Fig. 5B). With PEG 3350, the activity was lower than that observed in the pre-centrifugation control or when the pellet was resuspended in buffer with PEG 3350. By contrast, with citrate, the activity was similar to both control conditions (pre-centrifugation or when the pellet was resuspended in buffer and citrate). These data indicate that the phase transition of TIR is partially reversible in PEG 3350 and irreversible in sodium citrate. To confirm these findings, we centrifuged the resuspended samples to examine the fractions visually by SDS-PAGE. In the sample initially prepared with PEG 3350 and resuspended in buffer alone, a faint band was present in the supernatant fraction. However, this band was absent in the sample initially prepared with sodium citrate (Fig. 5C). These data confirm that the TIR phase transition is partially reversible in PEG 3350 and irreversible in sodium citrate, at least under these conditions.

Next, we evaluated the effect of pH on TIR precipitation and NADase activity. There was virtually no increase in TIR precipitation in the absence of PEG. However, in the presence of 25% PEG, TIR precipitation increased with increasing pH (Fig. 5D). Under these same conditions, no activity was apparent below pH 6. *K*_m_ remained constant (Fig. 5E) and *k*_cat_ increased from pH 6.5 to pH 8.5 (Fig. 5F). A corresponding increase in *k*_cat_/*K*_m_ was responsible for the increase in catalytic efficiency above pH 7 (Fig. 5G). Notably, the increase in *k*_cat_/*K*_m_ correlated with precipitation (Figs. 5D and 5G), again indicating that a phase transition increases TIR activity.

We performed negative stain electron microscopy to directly visualize TIR aggregation *in vitro*. Protein visualization is not possible with PEG because macroviscogens themselves are stained, confounding image analysis. Therefore, we performed this experiment with citrate. In the absence of citrate, we observed borderline fibrillar structures and protein aggregates, but overall, there were no consistent structures (Fig. 5H). However, in the presence of citrate, circular particles emerged (Fig. 5H). These results indicate that, *in vitro*, TIR aggregates into higher-order, ring-like structures.

### Physiologic stresses, both pathogen and non-pathogen, induce multimerization of TIR-1/SARM1 into visible puncta within intestinal epithelial cells

To visualize TIR-1 aggregation *in vivo*, we used CRISPR/Cas9 to insert wrmScarlet at the C-terminus of the endogenous *C. elegans tir-1* locus, which labeled all *tir-1* isoforms. In uninfected animals, TIR-1::wrmScarlet is barely detectable in intestinal epithelial cells (Fig. 6A). However, *P. aeruginosa* infection caused TIR-1::wrmScarlet to multimerize into visible puncta within intestinal epithelial cells (Fig. 6A and 6B). We distinguished TIR-1::wrmScarlet puncta from autofluorescent gut granules by comparing images in the red and green fluorescence channels. TIR-1::wrmScarlet puncta are those that are seen in the red, but not the green fluorescence channel (arrowhead in Fig. 6), as opposed to gut granules, which can be seen in both channels (asterisks in Fig. 6). Importantly, *tir-1(RNAi)* ablated TIR-1::wrmScarlet formation in the red fluorescence channel, but did not affect gut granule auto-fluorescence (Fig. 6A and 6B).

**Figure 6.**
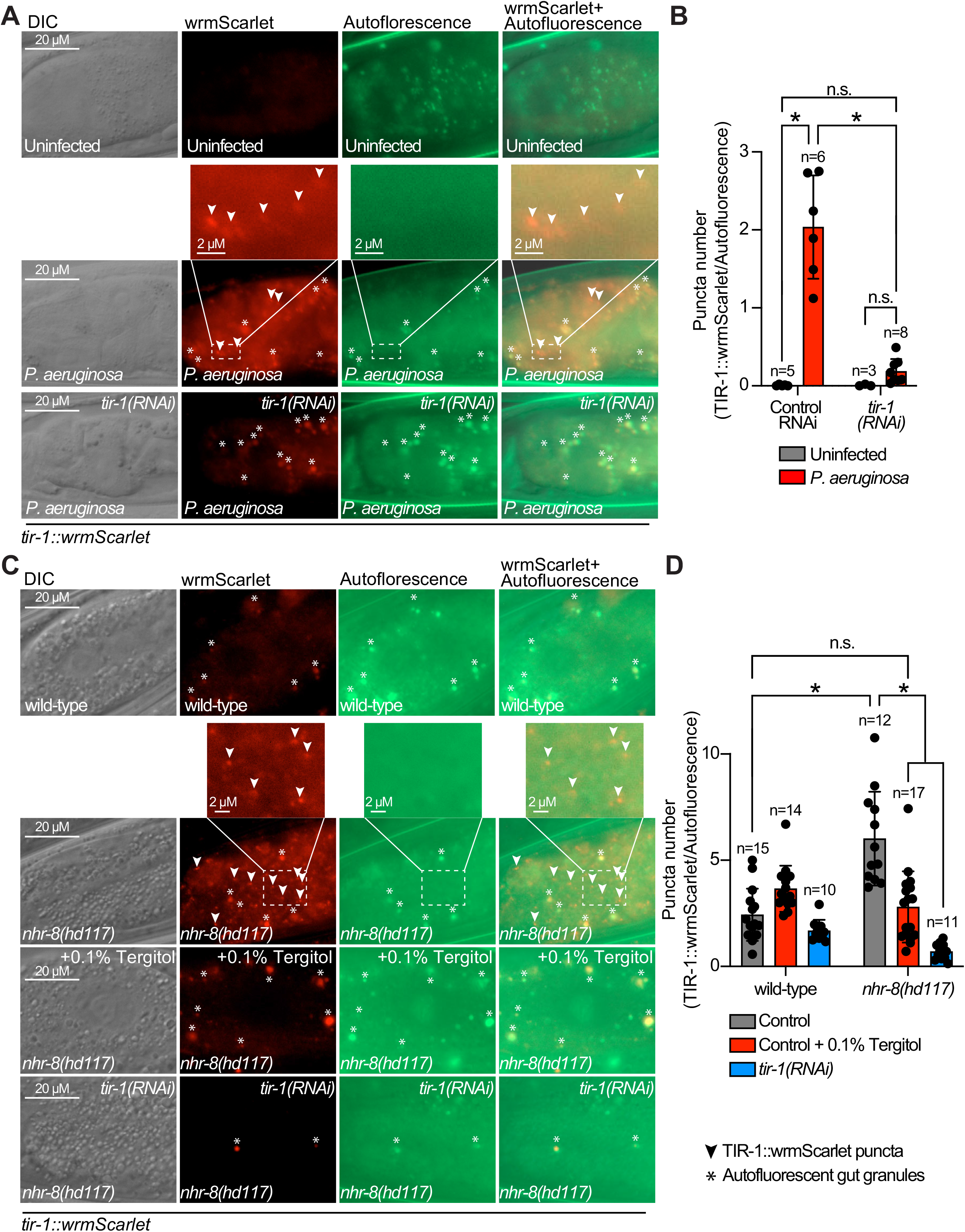
Physiologic stresses, both pathogen and non-pathogen, induce multimerization of TIR-1/SARM1 into visible puncta within intestinal epithelial cells. **(A)** Images of animals expressing TIR-1::wrmScarlet in the indicated conditions. All *tir-1*::*wrmScarlet* animals were treated with *glo-3(RNAi)* to deplete autofluorescent gut granules. Representative images for each condition are displayed. Red fluorescent channel images display both TIR-1::wrmScarlet fluorescence and autofluorescent signal, while the green fluorescent channel only displays signals from autofluorescent gut granules. TIR-1::wrmScarlet puncta are indicated by arrowheads and autofluorescent gut granules by asterisks. **(B)** The number of puncta present in the last posterior pair of intestinal epithelial cells in both the red (*tir-1*::*wrmScarlet*) and green (autofluorescence) fluorescent channels were quantified using Fiji image analysis software. Each data point is the ratio of red/green puncta from one animal. The n is indicated for each condition. **(C)** Images of *tir-1*::*wrmScarlet* and *nhr-8(hd117)*;*tir-1::wrmScarlet* animals as described in (A) exposed to either 0.1% Tergitol or *tir-1(RNAi)*. **(D)** Quantification of the ratio of red (TIR-1::wrmScarlet) to green (autofluorescence) puncta with indicated conditions as described in (B). Scale bar distance is indicated in all images. *equals p<0.05 by two-way ANOVA with Tukey multiple comparison testing.

Sterol depletion in *nhr-8(hd117)* animals also induced TIR-1::wrmScarlet puncta formation in intestinal epithelial cells (Fig. 6C and 6D). Importantly, TIR-1::wrmScarlet puncta formation in *nhr-8(hd117)* animals was entirely suppressed by solubilizing cholesterol in the growth media with Tergitol (Fig. 6C and 6D). Additionally, treatment of *nhr-8(hd117)* animals expressing TIR-1::wrmScarlet with *tir-1(RNAi)* inhibited fluorescence and puncta formation (Fig. 6C and 6D).

Taken together, these data demonstrate for the first time that physiological stressors, both pathogen and non-pathogen, induce TIR-1 multimerization into puncta within intestinal epithelial cells, which superactivates the intrinsic NADase activity of this protein complex to activate the p38 PMK-1 innate immune pathway.

### Sterol scarcity primes p38 PMK-1 immune defenses

Since *C. elegans* require cholesterol to survive bacterial infection and must obtain this essential metabolite from their diet, we hypothesized that, when environmental sterols are limited, activation of the p38 PMK-1 pathway represents an evolutionary adaptation that primes immune effector expression to anticipate challenges from bacterial pathogens. To test this hypothesis, we examined the expression of innate immune effector genes during bacterial infection in the presence and absence of cholesterol supplementation. Interestingly, the induction of *irg-4*p::*gfp* (Fig. 7A), *irg-5*p::*gfp* (Fig. 7B) and T24B8.5p::*gfp* (Fig. 7C) during *P. aeruginosa* infection was enhanced when nematodes were infected on media that did not contain supplemented cholesterol. Consistent with these data, *P. aeruginosa* infection also led to increased activation of *irg-4*p::*gfp* (Fig. 7D) and *irg-5*p::gfp (Fig. 7E) when *nhr-8* was depleted by RNAi. These data suggest that cholesterol scarcity primes p38 PMK-1 immune defenses for subsequent pathogen encounter.

**Figure 7.**
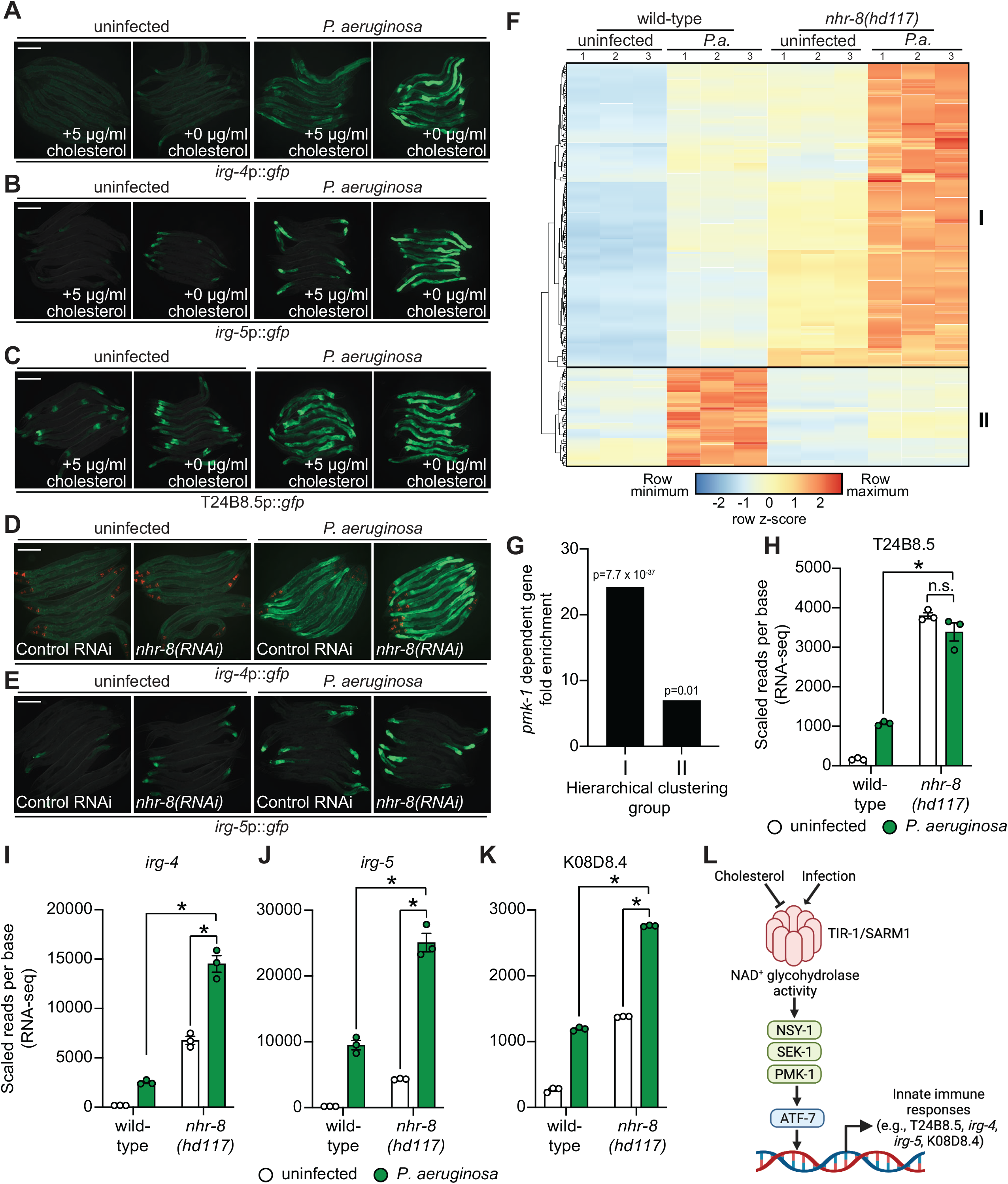
Sterol scarcity primes p38 PMK-1 immune defenses. **(A, B, C, D, E)** Images of the indicated transcriptional immune reporters under the indicated conditions. Scale bars in all images equal 200 μm. **(F)** A heat map compares the expression levels of the 184 genes that were both induced in wild-type animals during *P. aeruginosa* infection and differentially expressed (either induced or repressed) in *nhr-8(hd117)* mutants at baseline (each greater than 2-fold, q<0.05). To compare expression of these genes in wild-type and *nhr-8(hd117)* mutants, we scaled the expression level in each condition by calculating a z-score for each row and performed hierarchical clustering, which identified two main clusters (Cluster I contains 139 genes and Cluster II contains 45 genes). See also Table S3. **(G)** Enrichment of p38 PMK-1-dependent genes in Cluster I and II genes is shown. **(H, I, J, K)** mRNA-seq data for the indicated genes from the experiment described in (F) showing scaled reads per base from three biological replicates. *equals q<0.05 from RNA-seq analysis. **(L)** Model of p38 PMK-1 pathway activation during sterol scarcity and pathogen infection.

To provide further support for this hypothesis, we analyzed the expression pattern of p38 PMK-1-dependent transcripts in *nhr-8(hd117)* mutant animals during *P. aeruginosa* infection. Of the 472 genes that were induced in wild-type animals during *P. aeruginosa* infection, 184 were also differentially regulated (either induced or repressed) in *nhr-8(hd117)* animals that were infected with *P. aeruginosa*. To perform this analysis in an unbiased manner, we scaled the expression level in each condition for each of these 184 genes by calculating a row z-score and performed hierarchical clustering (Fig. 7F). We observed that these 184 genes group into two clusters: Cluster I had 139 genes and Cluster II contained 45 genes (Table S3). Cluster I was comprised mostly of genes whose expression in the absence of infection was higher in *nhr-8(hd117)* mutants than wild-type animals (129 of 139 genes). In addition, the majority of Cluster I genes were more strongly induced during *P. aeruginosa* infection in *nhr-8(hd117)* animals than wild-type animals. Cluster II, on the other hand, contained genes whose induction on *P. aeruginosa* were dependent on *nhr-8*. Interestingly, p38 PMK-1-dependent genes were strongly enriched among Cluster I, but not Cluster II, genes (Fig. 7G). Thirty-three of the 139 genes in Cluster I are known targets of the p38 PMK-1 immune pathway (24.17-fold enriched, hypergeometric p-value = 7.68 x 10^-37^), a group that includes the immune effectors T24B8.5 (Fig. 7H), *irg-4* (Fig. 7I), *irg-5* (Fig. 7J) and K08D8.4 (Fig. 7K). By contrast, only three genes in Cluster II are p38 PMK-1 dependent transcripts (6.95-fold enriched, hypergeometric p-value = 0.01) (Figs. 7F and 7G).

These data demonstrate that sterol scarcity primes the induction of innate immune effectors by activating the p38 PMK-1 immune pathway. Subsequent challenge by *P. aeruginosa* further drives immune activation in a manner that promotes clearance of pathogenic bacteria from the intestine (Fig. 1O).

## DISCUSSION

In this manuscript, we report two conceptual advances. First, we show that a phase transition of *C. elegans* TIR-1, an NAD^+^ glycohydrolase homologous to mammalian SARM1, underlies p38 PMK-1 immune pathway activation in the *C. elegans* intestine (Fig. 7L). Importantly, we demonstrate that physiologic stresses, both pathogen and non-pathogen, induce multimerization of TIR-1/SARM1 into visible puncta within intestinal epithelial cells. *In vitro* biochemical studies of purified protein and *C. elegans* genetic analysis revealed that multimerization and a phase transition of TIR-1/SARM1 engages its NAD^+^ glycohydrolase activity to activate the p38 PMK-1 innate immune pathway and provide protection during bacterial infection.

Second, we show that the TIR-1/SARM1 phase transition is modified by dietary cholesterol, revealing a new adaptive response that allows a metazoan host to anticipate environmental threats during micronutrient deprivation, a time of relative susceptibility to infection. *C. elegans* lack the ability to synthesize cholesterol *de novo* and must acquire dietary sterols from its environment to support development, fecundity, lifespan, and resistance against pathogen infection [19–25]. *C. elegans* larvae that encounter sterol-scarce environments are primed to enter an alternative developmental program, called dauer diapause, which promotes animal survival, in part, by altering their metabolism and growth to conserve energy [26–30]. Interestingly, only a fraction of sterol-starved *C. elegans* larvae enter dauer diapause and the remaining animals develop to adulthood [28, 34]. Here, we show that cholesterol starvation promotes multimerization of TIR-1/SARM1 into puncta, which activates the p38 PMK-1 immune pathway. Importantly, anticipatory activation of innate immune defenses in this manner during conditions of low cholesterol availability augments immune effector induction and promotes pathogen clearance from the intestine during a subsequent bacterial infection.

It is important to note that the conclusion of our manuscript regarding the role of cholesterol in *C. elegans* pathogen resistance is fundamentally different from the findings of Otarigho et al. [24]. These authors found that supraphysiologic concentrations of cholesterol given in the absence of infection drove the activation of immune effector genes in an RNA-seq experiment, five of which were dependent on *nhr-8* for their induction in this context using qRT-PCR. Together with genetic studies during pathogen infection, the authors conclude that NHR-8 controls immune genes downstream of multiple *C. elegans* pathways, including the p38 PMK-1 pathway. Indeed, in our genome-wide transcriptome profiling experiment, we observed that a small number of genes required *nhr-8* for their full induction during *P. aeruginosa* infection (Class II genes, Fig. 7F), a group that includes some genes downstream of the p38 PMK-1 pathway (Fig. 7G). However, we discovered that neither immune effectors, nor p38 PMK-1 targets, were enriched among the genes that were transcriptionally repressed upon exposure to 0 mg/ml cholesterol or in two *nhr-8* loss-of-function mutants (Figs. 1D, 1E, 2A, 2D, S1A and S2A). Rather, genes dependent on p38 PMK-1 for their transcription were strongly activated during cholesterol deprivation (both in wild-type animals exposed to 0 mg/mL cholesterol and in two *nhr-8* mutants at baseline, both in the absence of infection) (Figs. 1D, 1E, 2A, 2D, S1A and S2A). In addition, the induction of these p38 PMK-1-dependent immune effectors were strongly augmented during pathogen infection (Class I genes, Figs. 7F and 7G). We confirmed in western blot experiments and genetic epistasis analyses that the p38 PMK-1 pathway is activated in multiple *nhr-8* loss-of-function mutants and in wild-type animals grown in the absence of exogenous cholesterol (Fig. 2). We also employed two orthologous assays to demonstrate that the hypersusceptibility of multiple *nhr-8* mutants to *P. aeruginosa* infection is due to sterol deficiency in these animals (Figs. 1M and 1N). Finally, we show that low cholesterol in *nhr-8* mutants leads to markedly enhanced induction of p38 PMK-1-dependent immune effector expression during *P. aeruginosa* infection (Class I genes, Fig. 7F) in a manner that promotes clearance of pathogen from the intestine (Fig. 1O). Thus, our study reveals that cholesterol deficiency primes protective activation of the p38 PMK-1 immune pathway in the *C. elegans* intestine.

We previously demonstrated that a *C. elegans* nuclear hormone receptor, NHR-86, a member of a family of ligand-gated transcription factors, surveys the chemical environment to activate the expression of immune effectors [50]. Interestingly, NHR-86 targets immune effectors whose basal regulation requires the p38 PMK-1 immune pathway. However, NHR-86 functions independently of PMK-1 and modulates the transcription of these infection response genes directly. NHR-86 is a nematode homolog of the mammalian nuclear hormone receptor hepatocyte nuclear factor 4 (HNF4), a family of NHRs that expanded dramatically in *C. elegans* (259 HNF4 homologs are encoded in the *C. elegans* genome) [51, 52]. One potentially unifying hypothesis is that HNF4 homologs detect pathogen- or host-derived ligands that are associated with infection and activate pathogen-specific immune defenses. In this model, the p38 PMK-1 pathway functions as a rheostat that receives inputs from signals associated with potentially dangerous environmental conditions to prime host immune effector genes. Indeed, inputs from chemosensory neurons, bacterial density, tissue damage and nucleotide metabolism also regulate the tonic level of p38 PMK-1 pathway activity [3, 53–57], suggesting that p38 PMK-1 phosphorylation is adjusted to anticipate dangerous threats during periods of vulnerability. Thus, upon encountering a pathogen, immune defenses can be further augmented by other mechanisms that detect pathogen specifically.

SARM1, the mammalian homolog of TIR-1, has a well-characterized role in promoting Wallerian degeneration, a form of neuronal degeneration induced by axon injury [58, 59]. In *in vitro* models of axon injury, SARM1 NADase activity depletes NAD^+^ and induces axonal death [58]. Loring et al. reported that a phase transition of mammalian SARM1 potentiates its intrinsic NADase activity, as we observed for *C. elegans* TIR-1 [15]. These authors found that treatment with high concentrations of citrate, a precipitant that induces protein aggregation, caused *C. elegans* TIR-1/SARM1 to form puncta in neurons. Exposure of *C. elegans* to citrate was also correlated with enhanced axonal degeneration. Citrate exposure in this experiment does not represent a physiologically relevant stress, however. Thus, the findings reported here that TIR-1/SARM1 oligomerizes in response to a pathogen and non-pathogen stress provides an important characterization of the mechanism inferred by Loring, et al, and demonstrates that this phenomenon occurs in non-neuronal tissues in the regulation of intestinal immunity. Interestingly, the enzyme kinetics of TIR-1 are strikingly similar to mammalian SARM1 – the activity of both proteins is exquisitely dependent on precipitation. Thus, the mechanism of TIR-1/SARM1 activation is conserved across millions of years of evolution, functions in different cellular contexts, and underlies both axonal degeneration following neuronal injury and the regulation of the p38 PMK-1 pathway activation in the control of intestinal immunity.

## ACKNOWLEDGEMENTS AND FUNDING SOURCES

The authors thank Alexandra Byrne, Victoria Czech, Lauren O’Connor, and Heather Loring for helpful discussions. The authors are also grateful to Melanie Trombly and Merin MacDonald for critical reading of the manuscript. This research was supported by R01 AI130289 (to R.P.W.), R21 AI163430 (to R.P.W.), an Innovator Award from the Kenneth Rainin Foundation (to R.P.W.), the Dan and Diane Riccio Fund for Neuroscience (to R.P.W. and P.R.T.), F30 AI150127 (to N.D.P.), T32 AI132152 (to N.D.P. and J.D.I.), T32 GM107000 (to N.D.P.), and R35 GM118112 (to P.R.T.). Some strains were provided by the *Caenorhabditis* Genetics Center, which is funded by the NIH Office of Research Infrastructure Programs (P40 OD010440).

## MATERIALS AND METHODS

### *C. elegans* and bacterial strains

The previously published *C. elegans* strains used in this study were: N2 Bristol [31], KU25 *pmk-1(km25)* [60], AU3 *nsy-1(ag3)* [1], ZD101 *tir-1(qd4)* [32], AA968 *nhr-8(hd117)* [34], AE501 *nhr-8(ok186)* [35], AU78 *agIs219* [T24B8.5p::*gfp*::*unc-54*-3’UTR; *ttx-3*p::*gfp*::*unc-54*-3’UTR] [32], AU306 *agIs44* [*irg-4*p::*gfp*::*unc-54*-3’UTR*; myo-2*p:*:mCherry*] [61], AY101 *acIs101* [p*DB09.1(irg-5*p::*gfp)*; pRF4(*rol-6(su1006)*)] [33], TJ356 *zIs356* [*daf-16*p::*daf-16a/b*::*gfp* + pRF4(*rol-6(su1006)*)], *MGH167 sid-1(qt9); alxIs9[vha-6p::sid-1::SL2::GFP]*. The strains developed in this study were: RPW278 *nhr-8(hd117)*;*agIs219*, RPW317 *tir-1(qd4)*;*nhr-8(hd117)*, RPW339 *tir-1(ums47[E788A])*;*agIs219*, RPW369 *tir-1(ums54[ΔSAM])*;*agIs219*, RPW374 *tir-1(ums55[G747P])*;*agIs219*, RPW381 *tir-1(ums56[H833A])*;*agIs219*, RPW386 *tir-1(ums57[tir-1::3xFLAG])*;*agIs219*, RPW387 *tir-1(ums58[tir-1[E788A]::3xFLAG])*;*agIs219*, RPW388 *tir-1 (ums59[tir-1[ΔSAM]::3xFLAG])*;*agIs219*, RPW 389 *tir-1(ums60[tir-1[G747P]::3xFLAG])*;*agIs219*, RPW403 *tir-1(ums63[tir-1::wrmScarlet])*, RPW404 *nhr-8(hd117)*;*tir-1(ums63[tir-1::wrmScarlet])*. Bacteria used in this study are *Escherichia coli* OP50, *E. coli* DH5α, *E. coli* HT115(DE3), and *Pseudomonas aeruginosa* strain PA14 [62].

### *C. elegans* growth conditions and lipid supplementation

*C. elegans* strains were maintained on standard nematode growth medium (NGM) plates [0.25% bacto peptone, 0.3% sodium chloride, 1.7% agar (Fisher), 5 μg/mL cholesterol (Sigma-Aldrich, BioReagent grade), 25 mM potassium phosphate pH 6.0, 1 mM magnesium sulfate, 1 mM calcium chloride] with *E. coli* OP50 as a food source, as described [31]. For low cholesterol medium (0 μg/mL cholesterol), NGM was prepared without cholesterol supplementation, while 0.1% ethanol was added to maintain an equivalent ethanol concentration. For high cholesterol medium, cholesterol was dissolved in ethanol at 20 mg/mL and added to NGM at a final concentration of 80 μg/mL immediately prior to pouring plates. For all assays with high cholesterol medium, NGM containing 0.4% ethanol and 5 μg/mL cholesterol were used as control plates. Cholesterol solubilization assays were performed by supplementing NGM containing 5 μg/mL cholesterol with either 0.1% Tergitol (Sigma-Aldrich) or 0.1% Triton X-100 (Sigma-Aldrich). For assays using media solidified with agarose, NGM plates were prepared with 1.7% Ultrapure agarose (ThermoFisher Scientific) in place of agar. All fatty acids were purchased from Nu-Check-Prep Inc. and supplementation performed as previously described with modification [63, 64]. Fatty acids were dissolved in 50% ethanol and added at a final concentration of 1 mM to NGM agarose containing 0 μg/mL cholesterol and 0.1% Tergitol immediately prior to plate pouring. Prior to all assays, plates supplemented with lipids and control plates were seeded with *E. coli* OP50 and grown for 24 hours at room temperature. Assays were performed by picking 10-20 gravid adult animals to either lipid-supplemented or matched control plates. Animals were maintained on the plates for 14 hours at 20°C after which they were removed. Eggs laid on the plate were allowed to hatch and develop to the L4 stage at 20°C. For low cholesterol assays, animals were grown for two generations on NGM containing 0 μg/mL cholesterol.

### *C. elegans* strain construction

CRISPR/Cas9 was used to generate *tir-1* mutants in both wild-type and TIR-1::3xFLAG backgrounds, as described [65, 66]. All CRISPR/Cas9 reagents were purchased from Integrated DNA Technologies. Target guide sequences were selected using the CHOPCHOP web tool [67]. ssODN and dsDNA repair templates contained indicated edits, deletions or insertions with 35 bp flanking homology arms. crRNA guide and ssODN sequences are listed in Table S4. For wrmScarlet dsDNA repair template, wrmScarlet was PCR amplified with 35 bp flanking homology arms. PCR was gel purified, diluted to 100 ng/μL, melted and reannealed using a thermal cycler (95 °C -2 minutes; 85 °C – 10 seconds, 75 °C – 10 seconds, 65 °C – 10 seconds, 55 °C – 10 seconds, 45 °C – 10 seconds, 35 °C – 10 seconds, 25 °C – 10 seconds, 4 °C – hold. Ramp down 1 °C per second), and used immediately for injection. A mixture of 0.25 μg/μL Cas9, 0.1 μg/μL tracrRNA and 0.056 μg/μL crRNA were incubated for 15 minutes at 37°C. 0.11 μg/μL ssODN or 25 ng/μL dsDNA and 40 ng/μL pRF4(*rol-6(su1006)*) plasmid were added to the mixture, centrifuged and microinjected into young adult animals carrying the *agIs219* transgene, *tir-1(ums57[tir-1::3xFLAG])*;*agIs219*, or N2. The F1 progeny were screened for Rol phenotypes 3-4 days after injection and then for indicated edits using PCR and Sanger sequencing. Primer sequences used for genotyping are listed in Table S4.

### Feeding RNAi

Knockdown of target genes was performed by feeding *C. elegans E. coli* HT115 expressing dsRNA targeting the gene of interest, as previously described with modification [68–70]. In brief, HT115 bacteria expressing dsRNA targeting genes of interest were grown in Lysogeny broth (LB) Lennox medium containing 50 μg/mL ampicillin and 15 μg/mL tetracycline overnight with shaking (250 rpm) at 37°C. Overnight cultures were seeded onto NGM containing 5 mM IPTG and 50 μg/mL carbenicillin and incubated at 37°C for 16 hours after which synchronized L1 animals were transferred to bacterial lawns and allowed to grow until the L4 stage.

### *C. elegans* Bacterial Infection and Colonization Assays

“Slow killing” *P. aeruginosa* infection experiments were performed as previously described [71, 72]. Wild-type is either N2 or *agIs219*. In brief, a single colony of *P. aeruginosa* PA14 was inoculated into 3-mL of LB medium and grown with shaking (250 rpm) at 37°C for 14 hours. 10-μL of overnight culture was spread onto 35-mm petri dishes containing 4-mL slow killing agar (0.35% peptone, 0.3% sodium chloride, 1.7% agar, 5 μg/mL cholesterol, 25 mM potassium phosphate pH 6.0, 1 mM magnesium sulfate, 1 mM calcium chloride). Plates were incubated for 24 hours at 37°C and for approximately 24 hours at 25°C. Immediately prior to starting the assay 0.1 mg/mL 5-fluorodeoxyuridine (FUDR) was added on top of the agar to prevent progeny from hatching. Animals used in all assays were grown at 20°C with specified growth conditions. For assays involving high cholesterol or nonionic detergents, slow killing agar plates were prepared with either 80 μg/mL cholesterol (Sigma-Aldrich), 0.1% Tergitol (Sigma-Aldrich), or 0.1% Triton X-100 (Sigma-Aldrich). For experiments using plates containing 80 μg/mL cholesterol, matched control plates containing the equivalent ethanol concentration (0.4%) and 5 μg/mL cholesterol were prepared. All pathogenesis and lifespan assays are representative of three biological replicates. Sample sizes, mean lifespan, and p values for all trials are shown in Table S2.

Colony forming units of *P. aeruginosa* were quantified in the intestine of *C. elegans* as previously described with modifications [72, 73]. Briefly, *C. elegans* animals were exposed to lawns of *P. aeruginosa*, which were prepared as previously described, for 24 hours. Animals were then picked to NGM plates lacking bacteria and incubated for 10 minutes to remove external *P. aeruginosa*. Animals were then transferred to a second NGM plate after which 10-11 animals per replicate were collected, washed with M9 buffer containing 25 mM tetramisole (Sigma-Aldrich) and 0.01% Triton X-100 (Sigma-Aldrich), and ground with 1.0 mm silicon carbide beads (BioSpec Products). *P. aeruginosa* CFUs were quantified from serial dilutions of the lysate grown on LB agar.

### Gene expression analysis and bioinformatics

2,000 synchronized L1 stage *C. elegans* of the indicated genotypes were grown to the L4 stage and harvested by washing with M9. For expression analysis of *C. elegans* genes during *P. aeruginosa* infection, animals at the L4 stage animals were transferred by washing to plates containing *E. coli* OP50 or *P. aeruginosa* PA14 lawns. Animals were exposed for four hours and subsequently harvested by washing with M9. RNA was isolated using TriReagent (Sigma-Aldrich), column purified (Qiagen), and analyzed by 100 bp paired-end mRNA-sequencing using the BGISEQ-500 platform (BGIAmericasCorp) with >20 million reads per sample. Raw fastq reads were evaluated by FastQC (version 0.11.5), clean reads were aligned to the *C. elegans* reference genome (WBcel235) and quantified using Kallisto (version 0.45.0) [74]. Differentially expressed genes were identified using Sleuth (version 0.30.0) [75]. For identification of differentially expressed genes from animals grown in the absence (0 μg/mL) versus presence (5 μg/mL) of cholesterol supplementation, we analyzed previously generated publicly available RNA-seq data from Otarigho et al. [24] using eVITTA [76]. Pearson correlation statistical analysis was performed using Prism 9.0. Innate immune genes were identified using DAVID Bioinformatics database biological process gene ontology (GO) term innate immune response [77]. Heat-maps of differentially expressed genes were generated using pheatmap (version 1.0.12). Gene set enrichment analysis of RNA-seq was performed using GSEA (version 4.1.0) [78] with a custom gene set database containing p38 PMK-1 dependent genes generated from previously published RNA-seq of uninfected *pmk-1(km25)* animals [79]. Differential gene expression was defined as a fold change (FC) versus wild-type greater than 2 and q less than 0.01.

For the qRT-PCR studies, RNA was reverse transcribed to cDNA using the iScript^TM^ cDNA Synthesis Kit (Bio-Rad Laboratories, Inc.), amplified and detected using Syber Green (Bio-Rad Laboratories, Inc.) and a CFX384 machine (Bio-Rad Laboratories, Inc.). The sequences of primers that were designed for this study are presented in Table S4. Other primers were previously published [4, 80–82]. All values were normalized against the geometric mean of the control genes *snb-1* and *act-3*. Fold change was calculated using the Pfaffl method [83].

### Immunoblot analyses

Protein lysates from 2,000 *C. elegans* grown to the L4 larval stage on *E. coli* OP50 on NGM agar were prepared as previously described with modification [50, 84]. Harvested animals were washed twice with M9 buffer and resuspended in RIPA Buffer (Cell Signaling Technology, Inc.) containing 1x Halt Protease and Phosphatase inhibitor (ThermoFisher Scientific). Samples were lysed using a teflon homogenizer, centrifuged, and protein was quantified from the supernatant of each sample using the DC protein assay (Bio-Rad Laboratories, Inc.). NuPAGE^TM^ LDS sample buffer (ThermoFisher Scientific) was added to a concentration of 1X and 12.5-30 μg of total protein from each sample was resolved on NuPAGE^TM^ 4-12% BisTris (Phospho-PMK-1 and Total-PMK-1) or NuPAGE^TM^ 3-8% TrisAcetate (TIR-1::3xFLAG) gels (ThermoFisher Scientific), transferred to nitrocellulose membranes using a Trans-Blot Turbo Transfer System (Bio-Rad Laboratories, Inc.), blocked with 5% milk powder in TBST and probed with a 1:1000 dilution of an antibody that recognizes the doubly-phosphorylated TGY motif of PMK-1 (Cell Signaling Technology, #9211), a previously characterized total PMK-1 antibody [50], a monoclonal mouse anti-FLAG antibody (Sigma-Aldrich, M2), or a monoclonal mouse anti-tubulin antibody (Sigma-Aldrich, Clone B-5-1-2). Horseradish peroxidase (HRP)-conjugated anti-rabbit (Cell Signaling Technology, #7074) and anti-mouse IgG secondary antibodies (Abcam, #ab6789) were diluted 1:10,000 and used to detect the primary antibodies following the addition of ECL reagents (Thermo Fisher Scientific, Inc.), which were visualized using a BioRad ChemiDoc MP Imaging System. The band intensities were quantified using ImageJ version 2.0.0, and the ratio of active phosphorylated PMK-1 to total PMK-1 was calculated.

### TIR-1::wrmScarlet puncta visualization and quantification

For TIR-1::wrmScarlet puncta visualization, all TIR-1::wrmScarlet expressing animals were depleted of autofluorescent gut granules by exposure to *glo-3(RNAi)* from the L1 stage prior to all experiments. *P. aeruginosa* was prepared as described for bacterial infection and colonization assays. L4 animals expressing TIR-1::wrmScarlet were transferred to either OP50 or *P. aeruginosa* containing plates for 24 hours. For imaging, animals were transferred to 2% agarose pads, paralyzed with 300 mM sodium azide, and imaged with a 63x oil immersion lens. For Tergitol experiments, animals were grown on RNAi plates containing 0.1% Tergitol and the respective RNAi strains and imaged at the L4 stage. Image processing and analysis was performed using Fiji image analysis software. In each panel, the exposure time and processing for all images were identical. To quantify TIR-1::wrmScarlet puncta, maximum pixel intensity in the last posterior pair of intestinal epithelial cells in both the red (TIR-1::wrmScarlet) and green (autofluorescence) channels were quantified.

### Microscopy

Nematodes were mounted onto agar pads, paralyzed with 10 mM levamisole (Sigma) or 300 mM sodium azide and photographed using a Zeiss AXIO Imager Z2 microscope with a Zeiss Axiocam 506mono camera and Zen 2.3 (Zeiss) software.

### TIR-1 TIR domain expression and purification

The recombinant *C. elegans* TIR-1 TIR domain (TIR) was expressed in bacteria as previously described [85]. Briefly, the TIR domain cloned into the pET-30a(+) vector was transformed into chemically competent *E. coli* BL21(DE3) cells and maintained as a glycerol stock at −80°C. An inoculation loop was used to transfer the transformed bacteria into 5 mL of LB media with 50 µg/mL (final concentration) of kanamycin and the culture was grown overnight at 37°C while rotating. The next day, the cultures were diluted 1:400 in LB media with 50 µg/mL (final concentration) of kanamycin and grown at 37°C while shaking at 215 rpm until an OD_600_ of 0.7-0.8 was reached. After cooling, 50 µM IPTG (final concentration) was added to the culture to induce protein expression. The incubator temperature was decreased to 16°C and cells were incubated for an additional 16-18 h. Bacterial cells were collected by centrifugation at 3,000 *x g* for 15 min at 4°C, flash frozen in liquid nitrogen, and stored at −80°C until purification.

For purification, bacterial pellets were thawed on ice and then resuspended in Lysis Buffer (50 mM Tris•HCl pH 7.0, 300 mM NaCl, 10% (w/v) glycerol, 0.001% Tween 20) with Pierce^TM^ EDTA-free protease inhibitor mini tablets (ThermoFisher Scientific). The resuspension was incubated with 100 µg/mL lysozyme for 10 min at 4°C and sonicated with a Fisher Scientific Sonic Dismembrator sonicator (FB-705) in 50-mL batches at an amplitude of 30 for 20 s, pulsing for 1 sec on and 1 sec off, followed by a delay period of 20 s for a series of 12 cycles. Crude lysate was clarified at 21,000 *x g* for 25 min at 4°C, at which point the supernatant was applied to pre-equilibrated Strep-Tactin XT Superflow high-capacity resin (IBA Lifesciences) and allowed to enter the column by gravity flow; the Strep-Tactin resin had been equilibrated in Strep Wash Buffer (50 mM Tris•HCl pH 7.0, 300 mM NaCl). The column was washed with 30 column volumes of Strep Wash Buffer and the protein was eluted with 25 column volumes of Strep Elution Buffer (Strep Wash Buffer with 50 mM biotin). Protein eluted from the Strep-Tactin column was then applied to pre-equilibrated TALON Metal Affinity Resin (Takara) and allowed to enter the column by gravity flow; the TALON resin was equilibrated in His Wash 1 (50 mM Tris•HCl pH 7.0, 150 mM NaCl, 5 mM imidazole). A series of 15 column volume washes were applied (His Wash 1; His Wash 2: 50 mM Tris•HCl pH 7.0, 150 NaCl, 10 mM imidazole) and the protein was eluted in 20 column volumes of His Elution Buffer (50 mM Tris•HCl pH 7.0, 150 mM NaCl, 150 mM imidazole). The eluted protein was dialyzed overnight in Dialysis Buffer (50 mM Tris•HCl, pH 7.0, 150 mM NaCl). The next day, the protein was concentrated using a 10,000 NMWL Amicon Ultra-15 Centrifugal Filter Unit at 4°C and the protein concentration was determined by the Bradford assay. TIR was flash frozen in liquid nitrogen and stored at −80°C in 25-µL aliquots.

### Fluorescent NADase Assay

Nicotinamide 1,N^6^-ethenoadenine dinucleotide (ε-NAD, Sigma-Aldrich) is a fluorescent analog of NAD^+^ and was utilized in kinetic assays as a TIR-1 substrate. TIR-1 cleaves the nicotinamide moiety from ε-NAD to release nicotinamide and etheno-ADPR (ε-ADPR), which fluoresces (λ_ex_ = 330 nm, λ_em_ = 405 nm). Enzymatic activity was assayed in Assay Buffer (50 mM Tris pH 8.0, 150 mM NaCl; final concentration) using Corning® 96–well Half Area Black Flat Bottom Polystyrene NBS^TM^ Microplates for a final reaction volume of 60-μL, or Corning® 384-well Low Volume Black Round Bottom Polystyrene NBS^TM^ Microplates for a final volume of 20-µL; reactions were initiated by the addition of ε-NAD. ε-ADPR fluorescence intensity readings were taken in real time every 15 sec for 15-30 min using Wallac EnVision Manager Software and a PerkinElmer EnVision 2104 Multilabel Reader. Fluorescence intensity readings (λ_ex_ = 330 nm, λ_em_ = 405 nm) were converted to [ε-ADPR] with an ε-ADPR standard curve, which was produced by incubating fixed concentrations (0-400 µM) of ε-NAD with excess ADP-ribosyl cyclase and plotting the peak fluorescence intensity values against [ε-ADPR]. The activity was linear with respect to time under all conditions tested.

### Effect of crowding agents and sodium citrate on the activity of the TIR domain

The enzymatic activity of the TIR domain was evaluated using the Fluorescent Assay described above. First, the enzyme concentration dependence was determined. For purified protein, the enzyme (0-32.5 µM; final concentration) was added to Assay Buffer in duplicate, briefly incubated at room temperature for 10 min, and the reaction was initiated with 1 mM ε-NAD. Fluorescence intensity was monitored in real time every 15 sec for 15 min.

To evaluate the effect of crowding agents and sodium citrate on TIR activity, stock solutions of the additives were made. 50% (w/v) solutions of PEGs 8000, 3500, 1500, and 400, as well as 60% (w/v) solutions of dextran, sucrose, and glycerol were prepared and filtered. Initially, the concentration dependence of TIR in the presence of PEG 3350 was determined. TIR (0-15 µM; final concentration) was added to Assay Buffer with 25% PEG 3350 (final concentration) in duplicate, incubated for 10 min at room temperature, initiated with 1 mM ε-NAD, and monitored every 15 sec for 15 min. The concentration dependence of TIR in 25% PEG 3350 was used to establish that 2.5 µM TIR could be used to enable robust kinetic analyses. Next, the effect of viscogens on TIR activity was evaluated by adding 2.5 μM TIR (final concentration) in duplicate to Assay Buffer with or without 25% w/v of the viscogens (final concentration). Following a brief 10 min incubation period, the reaction was initiated with 1 mM ε-NAD and monitored every 15 sec for 20 min. The dose response of 2.5 μM TIR to the viscogens (0, 10, 20, and 30% final concentrations) was evaluated in the same manner and monitored every 15 sec for 15 min.

Sodium citrate was prepared as a stock solution of 2 M and filtered. The concentration dependence of TIR in the presence of 500 mM sodium citrate was determined. TIR (0-15 µM; final concentration) was added to Assay Buffer with 500 mM citrate (final concentration) in duplicate, incubated for 10 min at room temperature, and then the reaction was initiated with 1 mM ε-NAD and monitored every 15 sec for 15 min. Additionally, the dose dependence of sodium citrate was determined in duplicate. 2.5 µM TIR was added to Assay Buffer with (0-1000 mM) sodium citrate and briefly incubated at room temperature for 10 min. The reaction was initiated with 1 mM Ε-NAD, and fluorescence was monitored every 15 sec for 15 min.

In all cases, the fluorescence intensity was converted to ε-ADPR using the ε-ADPR standard curve described above. Slopes of the progress curves yielded the velocities of the reactions, which were plotted in GraphPad Prism.

### Effect of PEG 3350 and sodium citrate on steady state kinetics

Steady state kinetic reactions were carried out in Assay buffer with either PEG 3350 (0-25%; final concentration) or sodium citrate (0-1000 mM; final concentration); a constant concentration of 2.5 µM TIR was used in these assays. Reaction components were mixed in duplicate and incubated at room temperature for 10 min before initiating the reaction with ε-NAD (0-4000 µM, final concentration). Fluorescence intensity was monitored every 15 sec for 15 min. Using the ε-ADPR standard curve, the fluorescence was converted to [ε-ADPR]. The velocity of the reactions was calculated from the slope of the progress curve at each ε-NAD concentration and plotted in GraphPad Prism. Kinetic parameters were determined by fitting these velocities to the Michaelis-Menten equation (Eq. 1) at each PEG 3350 or sodium citrate concentration. *K*_m_, *k*_cat_, and *k*_cat_/*K*_m_ values were plotted against PEG 3350 or sodium citrate concentration.

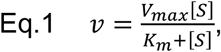

Where *V*_max_ is the maximum velocity, [*S*] is the substrate concentration, and *K*_m_ is the substrate concentration at half the maximum velocity.

### The TIR domain precipitates in PEG3350 and sodium citrate

5 µM TIR was incubated in Assay Buffer with PEG 3350 (0, 10, 17.5, and 25%; final concentration) or sodium citrate (0, 125, 250, 500, 750, and 1000 mM; final concentration) in duplicate at room temperature for 15 min. The precentrifugation control was removed and the remainder of the sample was centrifuged at 17,000 *x g* at 4 °C for 10 min. Following centrifugation, the supernatant was separated from the pellet and the pellet was resuspended in Assay Buffer with the respective concentration of PEG 3350 or sodium citrate. Samples were diluted 1:2 with gel loading buffer and run on an SDS-PAGE gel. Protein bands were stained by Coomassie and visualized on a BioRad Gel Doc EZ Gel Documentation System with Image Lab^TM^ Software. Representative images are shown.

Following resuspension of the pellet, all fractions were analyzed in the Fluorescent Assay. After the samples were aliquoted into the assay plate in duplicate, the enzymatic reaction was initiated with 1 mM ε-NAD and monitored every 15 sec for 15 min. Fluorescence intensity was converted to ε-ADPR concentration using the ε-ADPR standard curve. Slopes of the progress curves yielded the velocities of the reactions, which were plotted in GraphPad Prism.

### Effect of 1,6-hexanediol on TIR domain activity and precipitation

To evaluate the effect of 1,6-hexanediol on TIR activity, the Fluorescent Assay was performed in triplicate in the absence and presence of PEG 3350 or sodium citrate. For pure protein, 35 µM TIR (final concentration) was incubated in Assay Buffer at room temperature for 10 min with or without 2% 1,6-hexanediol (final concentration). For PEG 3350 or citrate, 2.5 µM TIR was incubated in Assay Buffer with 25% PEG 3350 or 500 mM sodium citrate (final concentrations) for 10 min at room temperature with or without 2% 1,6-hexanediol. The reactions were initiated with 1 mM ε-NAD and monitored every 15 sec for 15 min. Using the ε-ADPR standard curve, fluorescence was converted to [ε-ADPR] and the reaction velocities (i.e., slopes of the progress curves) obtained. Velocities were normalized for enzyme concentration and these normalized velocities were plotted in GraphPad Prism.

To determine whether 1,6-hexanediol can alter TIR precipitation, 1,6-hexanediol was added either before or after TIR precipitation in duplicate. To assess whether 1,6-hexanediol disrupts TIR precipitation, TIR precipitates were formed first by incubating 10 μM TIR (final concentration) with 25% PEG3350 or 500 mM sodium citrate (final concentration) at room temperature for 15 min. 1,6-hexanediol (0, 1, or 2%; final concentrations) was added to the mixture and incubated at room temperature for an additional 10 min. The precentrifugation control was removed and the remaining mixture was centrifuged at 21,000 *x g* for 10 min at 4°C. Supernatant fractions were removed, and the pellet was resuspended in Assay Buffer with the respective additive and concentration of 1,6-hexanediol. All fractions were run on an SDS-PAGE and stained with Coomassie Blue, and gels were imaged on a BioRad Gel Doc EZ Gel Documentation System with Image Lab^TM^ Software. Next, we determined if 1,6-hexanediol could prevent TIR precipitation. 10 µM TIR was incubated in Assay Buffer with 1,6-hexanediol (0, 1, or 2%; final concentration) for 15 min at room temperature. 25% PEG 3350 or 500 mM sodium citrate was added to the TIR-buffer-hexanediol mixture and incubated further for 10 min. Controls were removed and the samples were centrifuged at 21,000 *x g* for 10 min at 4°C. As before, supernatant fractions were removed, and pellet fractions were resuspended in Assay Buffer with respective additives and hexanediol concentrations. Fractions were analyzed by SDS-PAGE/Coomassie staining; representative images are shown.

### Phase Transition Reversibility

To evaluate the reversibility of the TIR phase transition, 5 µM of TIR was mixed with Assay Buffer and 25% PEG 3350 or 500 mM sodium citrate in duplicate (final concentrations). Following a 15 min incubation period at room temperature, precentrifugation controls were removed and the sample remaining was centrifuged at 17,000 *x g* for 10 min at 4°C. Supernatant fractions were separated from the pellet, which was resuspended in either Assay Buffer alone or Assay Buffer with respective additive. All fractions were analyzed for enzymatic activity in the Fluorescent Assay. Briefly, the precentrifugation, supernatant, and pellet fractions were aliquoted into the assay plates in duplicate and the reaction was initiated with 1 mM ε-NAD. Fluorescence was converted to [ε-ADPR] concentration with the ε-ADPR curve to yield the progress curves. The velocity of the reactions was taken as the slope of the line and the velocities were plotted in GraphPad Prism.

To validate the kinetic data, 10 µM TIR (final concentration) was incubated in Assay Buffer with either 25% PEG 3350 or 500 mM sodium citrate (final concentration) for 15 min at room temperature; control samples were incubated in Assay Buffer only. Samples were centrifuged at 21,000 *x g* for 10 min at 4°C, after which the supernatant was separated from the pellet. The pellets from samples initially prepared with additives were resuspended in either Assay Buffer alone or Assay Buffer plus the respective additive; this was not necessary for the sample initially prepared without additive since the protein is primarily located in the supernatant in this case. After removing another control sample (Additive lanes on gel), the resuspended samples were centrifuged again at 21,000 *x g* for 10 min at 4°C. As before, the supernatant was removed, and the pellet was resuspended in Assay Buffer with the respective additive. All samples were analyzed for protein content on an SDS-PAGE gel and stained with Coomassie Blue.

### Effect of pH on TIR domain precipitation and kinetics

To determine the effect of pH on TIR precipitation, the experiments described above were carried out in duplicate at pH values from 4.5-9.0. Briefly, 5 µM TIR (final concentration) was mixed with Assay Buffer (50 mM buffer; 150 NaCl) with and without 25% PEG 3350 (final concentration); for pH 4.5-5, sodium acetate buffer was used; for pH 5.5-6.5, MES was used; for pH 7-9, Tris was used. The samples were incubated for 15 min at ambient temperature, at which point the samples were centrifuged at 21,000 *x g* for 10 min at 4°C. The supernatant was removed, and the pellet was resuspended in either buffer alone or buffer with 25% PEG 3350 (final concentration); the presence or absence of 25% PEG 3350 and the buffer identity of the resuspension solution corresponded to the initial sample preparation. 10-µL 1 M Tris, pH 6.8 was added to each sample to neutralize the buffer before running on an SDS-PAGE gel and staining with Coomassie blue. Images of the gels were obtained on the BioRad Gel Doc EZ Gel Documentation System with Image Lab^TM^ Software. Representative images are shown. ImageJ was used to quantify the bands, which were plotted in GraphPad Prism.

Steady state kinetic analyses were also performed at each pH in 25% PEG 3350. A constant concentration of 2.5 µM TIR was used in these assays. Reaction components were mixed in quadruplicate and incubated at room temperature for 10 min before initiating the reaction with 0-2000 µM of ε-NAD (final concentration). Fluorescence intensity was monitored every 15 sec for 15 min. Using the ε-ADPR standard curve, the fluorescence was converted to [ε-ADPR]. The velocity of the reactions was calculated from the slope of the progress curve at each ε-NAD concentration. Kinetic parameters were determined by fitting these velocities to the Michaelis-Menten equation (Eq. 1) at each pH. The log of *K*_m_, *k*_cat_, and *k*_cat_/*K*_m_ values were determined and plotted in GraphPad Prism.

### Negative Stain Electron Microscopy

Negative stain EM on TIR (270 µg/mL) was performed in Assay Buffer (50 mM Tris, pH 8; 150 mM NaCl) with or without 500 mM sodium citrate. Samples were applied to glow-discharged, carbon- and formvar-coated copper grids and allowed to sit for 1 min and 30s before blotting excess liquid away. 1% uranyl acetate was used to fix the samples before image. Samples were images on a FEI Tecnai Spirit 12 microscope. The diameter of particles in the samples with citrate were analyzed in ImageJ.

### TIR domain mutants

TIR domain mutants were made using PCR based methods. The pET30a+ TIR-1 TIR domain construct was used as a template (0.2-2 ng/µL; final concentration). The manufacturer’s protocol for iProof^TM^ High-Fidelity DNA Polymerase (Bio-Rad Laboratories, Inc.) was followed using iProof HF buffer supplemented with 3% DMSO. 50-µL reaction volumes were used in the following protocol: initial denaturation for 3 min at 98°C, denaturation for 45 s at 98°C, annealing for 1:30 min at 45-72 °C, extension for 6 min at 72°C, final extension for 10 min at 72°C. Denaturation, annealing, and extension steps were repeated 30 times. The following day, *Dpn*I was added to the PCR reactions and incubated at 37°C for 2 h to digest the template DNA. The digest was transformed into chemically competent *E. coli* XL1-Blue cells. Transformants were grown overnight in LB media with 50 µg/mL kanamycin and mini-prepped (Promega). Mutagenesis was validated by Sanger sequencing (Genewiz). The mutants were expressed and purified as described for WT TIR.

Steady state kinetic analyses of the mutants were carried out in Assay Buffer with 25% PEG 3350 or 500 mM sodium citrate with 2.5 µM of the mutants (final concentrations). Reaction components were mixed in triplicate and incubated at room temperature for 10 min before initiating the reaction with 0-2000 µM of ε-NAD for PEG 3350 or 0-4000 µM for citrate. Fluorescence intensity was monitored every 15 sec for 17.5 min. Using the ε-ADPR standard curve, the fluorescence was converted to [ε-ADPR]. The velocity of the reactions was calculated from the slope of the progress curve at each ε-NAD concentration and plotted in GraphPad Prism. Kinetic parameters were determined by fitting these velocities to the Michaelis-Menten equation (Eq. 1) for each mutant.

To evaluate the precipitation capacity of TIR mutants, 10 µM (WT, G747P, E788Q, and H833A) or 3 µM (WT, E788A) of the enzyme was mixed with Assay Buffer (50 mM Tris, pH 8.0; 150 NaCl) with and without 25% PEG 3350 or 500 mM sodium citrate (final concentrations). Precipitation of the TIR^E788A^ mutant was evaluated at 3 µM due to low yields of the protein; WT TIR at 3 µM was included as the proper control. The samples were incubated for 15 min at ambient temperature, at which point the samples were centrifuged at 21,000 *x g* for 10 min at 4°C. The supernatant was removed, and the pellet was resuspended buffer with the respective additive (final concentration). Samples were diluted 1:1 with water before running on an SDS-PAGE gel. Stain free images of the gels were obtained on the BioRad Gel Doc EZ Gel Documentation System with Image Lab^TM^ Software. Representative images are shown. ImageJ was used to quantify the bands, which were plotted in GraphPad Prism.

### Statistical Analyses

Differences in survival of *C. elegans* in the *P. aeruginosa* pathogenesis assays were determined with the log-rank test after survival curves were estimated for each group with the Kaplan-Meier method. OASIS 2 was used for these statistical analyses [86]. qRT-PCR studies, intestinal CFU quantification, western blot band intensity quantification, TIR NADase activity, and TIR protein precipitation are presented as the mean ± standard error of the mean. Statistical hypothesis testing was performed with Prism 9 (GraphPad Software) using methods indicated in the figure legends. Sample sizes, mean lifespan, and p-values for all trials are shown in Table S2.

### Data Availability

The mRNA-seq dataset is available from the NCBI Gene Expression Omnibus using the accession number GSE178572. All other data are available in the manuscript.

---

**Supplementary Figure 1. Cholesterol scarcity activates intestinal innate immune defenses. (A)** An mRNA-seq experiment as described in Fig. 1D, except using a different *nhr-8* mutant allele: *nhr-8(ok186)*. **(B)** Images of T24B8.5p::*gfp* animals as described in Fig. 1G. **(C)** Images of T24B8.5p::*gfp* on agarose media with 0.1% Tergitol in the presence of supplemented fatty acids, as indicated. See also Fig. 1.

**Supplementary Figure 2. Cholesterol scarcity activates the p38 PMK-1 innate immune pathway. (A)** Gene set enrichment analysis (GSEA) of p38 PMK-1 targets in the mRNA-seq experiment as described in Fig 2D, except using a different *nhr-8* mutant allele: *nhr-8(ok186)*. **(B)** Images of T24B8.5p::*gfp* animals, as described in Fig. 2G. **(C)** Images of the transgenic *C. elegans*, in which *gfp* has been fused to the DAF-16 protein under the indicated conditions. Scale bars in all images equal 200 μm. See also Fig. 2.

**Supplementary Figure 3. Activation of the p38 PMK-1 pathway by cholesterol deprivation requires TIR-1 oligomerization and NADase activity. (A)** qRT-PCR data of *irg-4* in wild-type and mutant animals as described in Fig. 3G. **(B)** Immunoblot analysis of lysates from the indicated genotypes probed with an antibody that recognizes the FLAG epitope. **(C)** Images of T24B8.5p::*gfp* immune reporter containing the indicated genotypes. Scale bars in all images equal 200 μm. See also Fig. 3.

**Supplementary Figure 4. (A)** Enzyme concentration dependence in presence and absence of either 25% PEG 3350 or 500 mM sodium citrate (n=2). **(B)** Dose dependence of TIR NADase activity on citrate concentration as described in Fig. 4C, except using citrate (n=2). **(C)** Steady-state kinetic analysis of TIR as described in Fig. 4D, except in the presence of citrate (n=2). **(D-F)** From the steady-state kinetic analysis performed in Fig 4C, *K*_m_ **(D)**, *k*_cat_ **(E)**, and *k*_cat_/*K*_m_ **(F)** were determined at each citrate concentration. **(G)** Effect of citrate on TIR aggregation and associated activity as described in Fig. 4H (n=2, representative images shown). **(H)** Steady state kinetic analysis of TIR mutants as described in Fig. 4I, except in the presence of citrate (n=3). **(I, J)** Precipitation of TIR mutants as described in Fig. 4J and 4K, except in the presence of citrate (n=4, representative images shown). See also Fig. 4.

**Supplementary Figure 5. (A)** SDS-PAGE analysis of TIR protein soluble (S) and pellet (P) fractions following TIR incubation first with 25% PEG 3350 or 500 mM citrate and subsequent addition of 2% 1,6-hexanediol (n=2, representative image shown). **(B)** SDS-PAGE analysis of TIR protein soluble (S) and pellet (P) fractions following TIR incubation first with 0 or 2% 1,6-hexanediol and subsequent addition of 25% PEG 3350 or 500 mM citrate (n=2, representative image shown). See also Fig. 5.

**Supplementary Table S1. Genes significantly differentially expressed in *nhr-8(hd117)* and *nhr-8(ok186)* mutants compared to wild-type and in uninfected wild-type animals in the absence (0 μg/mL) versus presence (5 μg/mL) of cholesterol supplementation in the RNA-seq experiments presented in Figs. 1D, 1E and S1A.**

**Supplementary Table S2. Sample sizes, mean lifespan, and p values for the *C. elegans* pathogenesis assays.**

**Supplementary Table S3. Genes in Cluster 1 and Cluster 2 of the heat map shown in Fig. 6F.**

**Supplementary Table S4. Primer, crRNA guide and ssODN sequences designed for this study.**

