## Supplementary figures and images for "A TIR-1/SARM1 phase transition underlies p38 immune pathway activation in the *C. elegans* intestine"

### Supplementary Figure 1

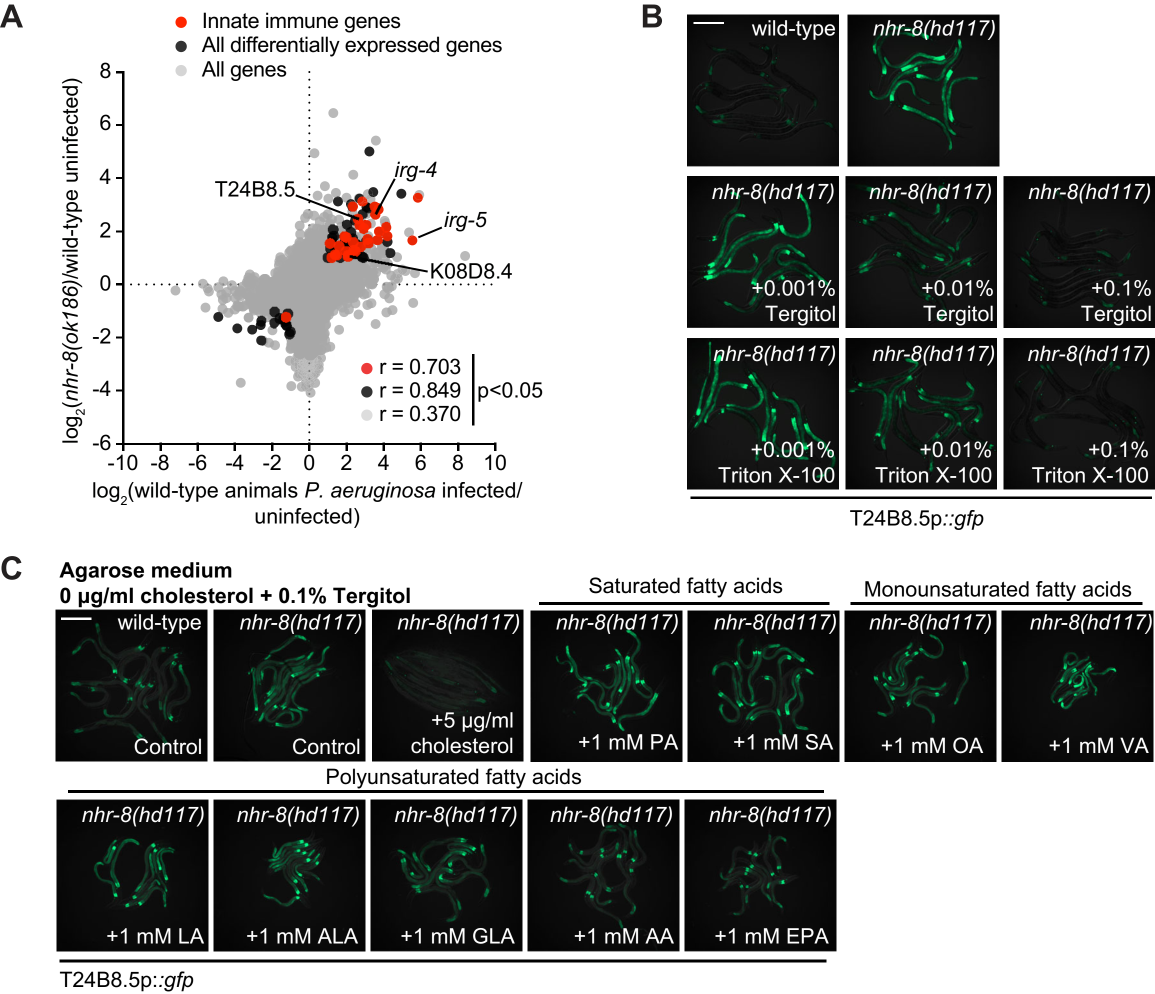

### Supplementary Figure 2

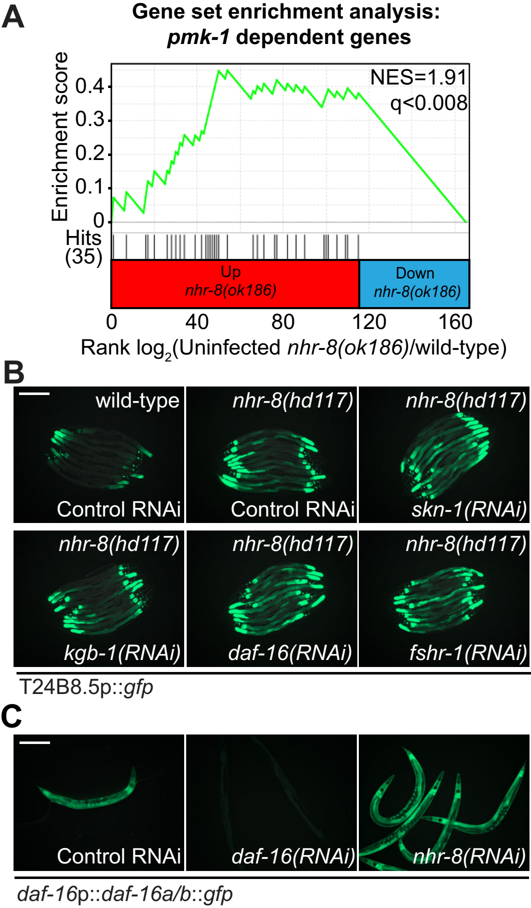

### Supplementary Figure 3

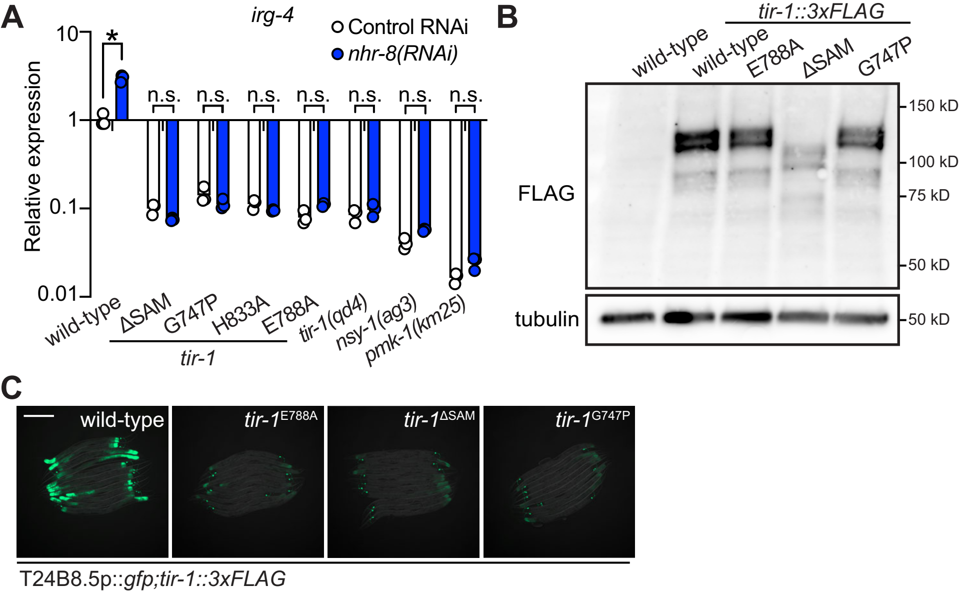

### Supplementary Figure 4

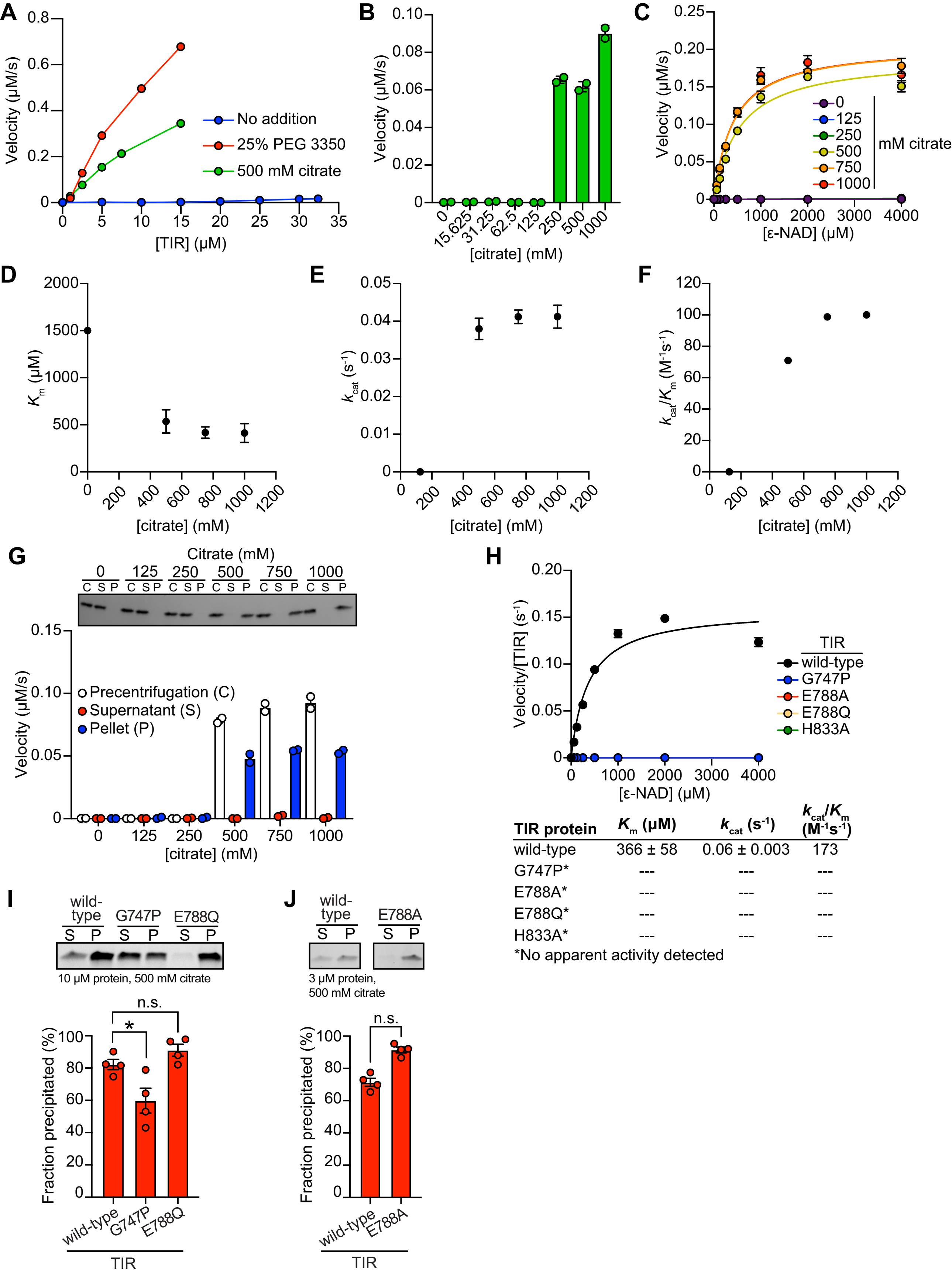


| TIR protein | $K_m$ (µM) | $k_{cat}$ ($s^{-1}$) | $k_{cat}/K_m$ ($M^{-1}s^{-1}$) |
|---|---|---|---|
| wild-type | 366 ± 58 | 0.06 ± 0.003 | 173 |
| G747P* | --- | --- | --- |
| E788A* | --- | --- | --- |
| E788Q* | --- | --- | --- |
| H833A* | --- | --- | --- |

*No apparent activity detected

### Supplementary Figure 5

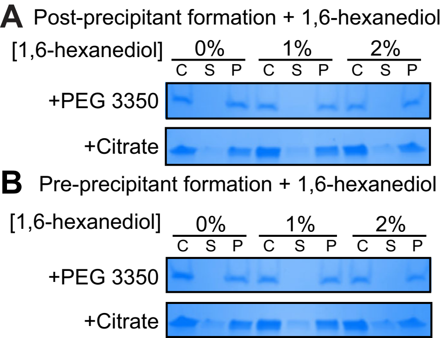
